# Preferred conformations in the disordered region of human CPEB3, a functional amyloid linked to memory consolidation

**DOI:** 10.1101/2020.05.12.091587

**Authors:** D. Ramírez de Mingo, D. Pantoja-Uceda, R. Hervás, M. Carrión-Vázquez, D. V. Laurents

## Abstract

While implicated in neurodegenerative diseases, amyloids are also essential to some physiological processes, including memory consolidation by neuronal-specific isoforms of the Cytoplasmic Polyadenylation Element Binding (CPEB) protein family. CPEB mediates memory persistence by the formation of self-sustaining amyloid assemblies through its intrinsically disordered region (IDR). Here, we characterize the atomic level conformation and ps-ns dynamics of the 426-residue IDR of human CPEB3 (hCPEB3), which has been associated with episodic memory in humans, by NMR spectroscopy. We found that the first 29 residues: M_1_QDDLLMDKSKTQPQPQQQQRQQQQPQP_29_, adopt a helical+disordered motif. Residues 86-93: P_83_QQPPPP_93_, and 166-175: P_166_PPPAPAPQP_175_ form polyproline II (PPII) helices. While the (VG)_5_ repeat motif is completely disordered, residues 200-250 adopt three partially populated α-helices. Residues 345–355, which comprise the nuclear localization signal (NLS), form a modestly populated α-helix and border a phosphoTyr which may mediate STAT5B binding. These findings allow us to suggest a model for nascent hCPEB3 structural transitions at single residue resolution, advancing that amyloid breaker residues, like proline, are a key difference between functional versus pathological amyloids. Besides revealing some aspects of the molecular basis of memory, these findings could aid the future development of treatments for post-traumatic stress disorder.

**Areas:** Biophysics, Structural Biology, Biochemistry & Neurosciences.

## Introduction

The molecular basis of long-term memory, which endures decades despite being built by ephemeral biomolecules, has long fascinated biochemists [1]. Seminal findings by Si, Lindquist and Kandel suggested that long-term changes in synaptic efficacy require a self-perpetuating amyloid state in the *Aplysia* CPEB (*Ap*CPEB) [2][3] . This change in CPEB’s conformation would lead to permanent alterations at the synapse constituting a physical substrate of memory storage. To attain the aggregated state necessary to stabilize memory, *Ap*CPEB contains a N-terminal IDR that is very rich in glutamine (Q) residues which losses α-helix and gains coiled-coil and β-sheet structure during amyloid formation *in* vitro [4][5].

The *Drosophila* homolog of CPEB, called Orb2, behaves in a similar fashion even though its N-terminal IDR has a lower glutamine residue content and a more tightly regulated amyloid formation [6][7][8][9]. Indeed, inhibition of Orb2 amyloid formation targeting the N-terminal IDR specifically impairs memory consolidation, but not short-term memory in *Drosophila* [6][10]. Due to numerous histidine (H) residues in the Orb2 Q/H-rich amyloid core comprised in the in the N-terminal IDR, pH may regulate this structure’s stability as suggested by CryoEM analysis [5] and characterization by NMR spectroscopy [11] In mammals, the N-terminal IDR of the neuronal-specific isoform of CPEB3 is crucial for amyloid formation and memory consolidation [12] [13]. The regulation of functional amyloid formation in mammalian CPEB3 appears to be even more sophisticated due to multiple mechanisms involving post-translational modifications [14], and feedback loops to maintain hCPEB3 expression levels [13]. Compared to the *Aplysia* and *Drosophila* homologs, hCPEB3’s content of glutamine residues in its 426-residues long IDR is lower and it contains diverse segments which are enriched for certain residues such as Ser, Ala, Pro, Gly+Val and hydrophobic residues (Table 1).

**Table 1:**
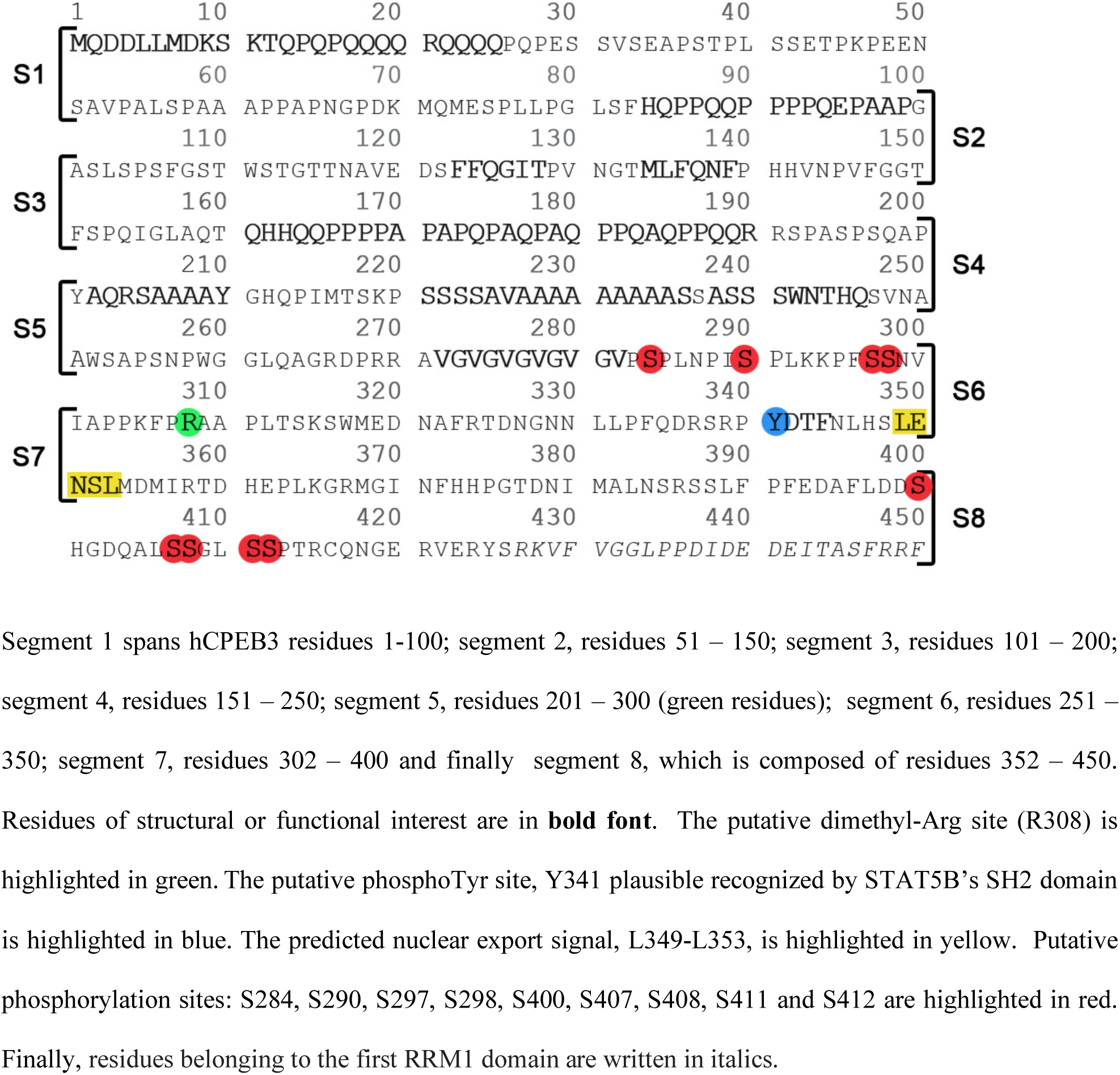
Sequence of the disordered N-terminal region of hCPEB3

Mammalian CPEB3 travels to distinct neuronal regions to carry out multiple functions, where the 426-residue long IDR plays a key role (**Figure 1A**). Following its synthesis, CPEB3 is SUMOlyated, which has been reported to block CPEB3 aggregation [14]. Upon neuronal stimulation, CPEB3, which is mostly cytoplasmic, travels to the nucleus. This process is mediated by the karyopherin IPO5 through interactions with the NLS in the first RNA recognition motif (RRM) of hCPEB3 [15]. Inside the nucleus, CPEB3 interacts with STAT5B, which normally activates the transcription of genes such as *EGFR*, triggering signaling cascades thought to promote memory consolidation [16][17][17]. CPEB3-STAT5B binding, driven by interactions between the IDR of hCPEB3 and residues 639–700 of STAT5B, downregulates STAT5B-dependent transcription [17]. However, the details of this interaction have not yet been addressed. By contrast, the 3D structure of the first CPEB3 RRM domain has been elucidated and revealed a β-hairpin (W471-G485) proposed to play a key role in RNA recognition [18]. This domain, as well as the second RRM domain and Zinc Finger (ZnF) motif, were reported to bind specifically to the 3’UTR mRNA of the AMPA receptor subunit GluR2 [19]. Together, CPEB3 and its target mRNA eventually exit the nucleus and can join distinct biomolecular condensates such as stress granules and neuronal granules, which provide physiological transport to dendritic spines, or to dendritic P-body-like granules [20], where CPEB3 stores and downregulates GluR2 mRNA translation [19].

**Figure 1.**
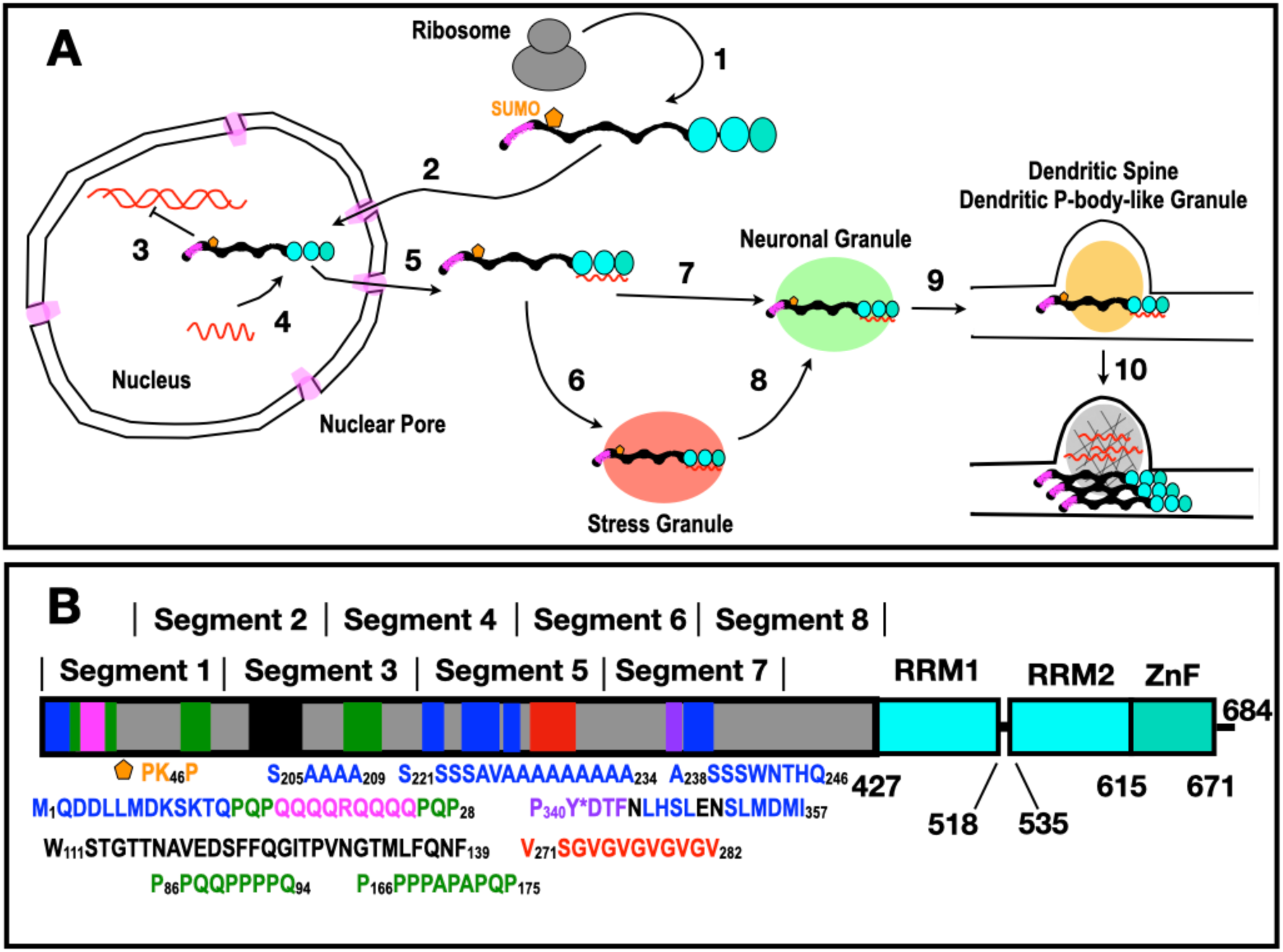
**Domain Structure and Activities of hCPEB3** A. hCPEB3 is present in multiple cellular compartments. Dendritic stimulation leads to temporary, phosphorylation-mediated short term memory and **1**. increased synthesis of the protein CPEB3. Composed of an N-terminal disordered region (**black**) which includes a Q-rich segment aiding functional aggregation (**magenta**), hCPEB3 also contains RRM domains (**cyan**) and a ZZ-Zinc finger domain (**turquoise**). **2**. Upon continued neuro-stimulation, CPEB3 enters the nucleus through the nuclear pore (light magenta), which is a macromolecular condensate. Once in the nucleus, CPEB3 indirectly regulates transcription through STAT5B **3,** and binds to certain mRNAs **4** (**red**). This binding suppresses translation. After exiting the nucleus through the nuclear pore, **6** CPEB3+mRNA may associate with a stress granule ( ) during moments of cellular stress. In the absence of stress (**7**) or its passing (**8**), CPEB3+mRNA will combine with another condensate called neuronal granules (**light green**) for transport to dendritic spines (**9**), where CPEB3+mRNA associate with still another class of condensate called a dendritic P-body-like structure (**golden**) (Cougot *et al.,* 2008) [103]. Further neuronal stimulation (**10**) causes synapse-specific deSUMOlyation, CPEB3 aggregation and translational activation of previously repressed mRNA, leading to morphological changes and fortification of the spine, which is proposed to be the basis of long-term memory. This is a simplified model based on (Kandel *et al.,* 2014) [104]. **B.** CPEB3 domain composition and its N-terminal intrinsically disordered domain (**gray**) contains key elements with preferred conformers colored **blue** for α-helix, **magenta** for polar amyloidogenic, **black** for hydrophobic amyloidogenic, **green** for PPII helix, **purple** for the putative phosphoTyr site, and **red** for highly disordered segments. The two RRM domain are colored **cyan** and the C-terminal Zinc Finger is shown in **turquoise**.

After synaptic activity in the hippocampus, SUMOylation of CPEB3 decreases [14] and CPEB3 converts, mediated by the IDR, from a translation repressor into a self-sustaining activator, promoting the translation of AMPA receptors [12]. This leads to structural modifications, including a more robust actin network, which fortify the spine and permanently enhance neurotransmission at a given particular synapse [13]. The hypothesis that CPEB functional amyloid formation is key for memory persistence in mammals is supported by the impairment in long-term memory and long-term potentiation in the CPEB3 conditional knockout mice [12]. In our species, the causal role of human CPEB3 (hCPEB3) in memory is corroborated by observations that persons carrying a rare CPEB3 allele, which leads to a decreased production of hCPEB3 protein, have episodic memory impairments [21].

The 426-residue long IDR of hCPEB3 plays a key role mediating memory persistence through this prion-like mechanism. It contains an amyloid-forming region spanning residues 1-200, and a condensate-promoting region formed by residues 250-426 which are linked by an alanine rich segment [22]. The full IDR is followed by two folded RRM which bind RNA and finally a ZZ-type ZnF domain (**Figure 1B**). Recent sequence and deletion mutational analyses of the IDR have begun to identify subregions key for aggregation, such as the first 30 residues [13]. However for the hCPEB3 IDR, programs to predict secondary structure tendencies give different outputs, Alpha Fold 2 structural predictions [23] are marked as low to very low confidence (see https://alphafold.ebi.ac.uk/entry/Q8NE35) and to date, no high-resolution experimental data on the partial structures or motions have been reported. Here, motivated by the key roles of the IDR in CPEB3 functions in transcription via association with STAT5B, as well as in CPEB prion-like aggregation required for memory persistence, we characterize the atomic level conformation and dynamics of the complete IDR of hCPEB3 by NMR spectroscopy, suggesting a speculative model for CPEB3 aggregation.

## Results

### hCPEB3’s IDR is chiefly disordered

As a first step to experimentally characterize hCPEB3’s IDR, we probed the complete 426 residue IDR of hCPEB3 by biophysical techniques and homonuclear NMR. Its fluorescence emission spectra, recorded at temperatures ranging from 2 to 70 °C, show emission maximum > 350 nm. This is consistent with its six Trp residues being solvent exposed and not buried in the hydrophobic core of a folded domain (**Sup. Fig. 1A**) [24]. The far UV CD spectrum of the hCPEB3 IDR also shows the hallmarks of a disordered protein, namely a minimum near 200 nm [25]. No spectral features indicative of α-helix and β-sheet; namely, minima at 208, 218 or 222 nm and no maximum at 195 nm, are evident (**Sup. Fig. 1B**). The 1D ^1^H and 2D ^1^H-^1^H NOESY spectra show ^1^H signals clustered into a narrow bands near the values observed for short, unstructured peptides (**Sup Fig. 1C**) [26][27]. The sequence alignment of several representative vertebrate CPEB3 proteins using the T-Coffee program is shown in **Sup. Fig. 2**. Very similar results were obtained from the Omega Clustal program (not shown). Whereas most IDPs show poor levels of sequence conservation, some stretches rich in hydrophobic residues, such residues M1-T12, W111-F139 and Y341-I357, are highly conserved. By contrast, glutamine rich, alanine rich and some proline rich segments are present only in mammals. Taking all these data together, the presence of large, stably folded domains in the IDR can be ruled out, but short segments with partly populated secondary structures could still be present.

### Atomic level characterization reveals partially structured elements in hCPEB’s N-terminal “disordered” region

To discover and characterize possible segments with partial secondary structure, we applied multidimensional heteronuclear NMR. As the full length IDR is too long to characterize by this methodology, we have followed the “divide and conquer” approach implemented by Zweckstetter *et al.* to characterize tau, a similarly sized IDP implicated in Alzheimer’s disease and other tauopathies [28]. As described in the **Material & Methods** section, and shown in **Table 1**, eight overlapping segments of 100 residues were characterized.

Using our powerful ^13^CO, ^15^N, ^1^HN based assignment strategy, over 99% of the main chain ^13^CO, ^13^Cα, ^15^N, ^1^HN and the ^13^Cβ resonances were assigned for residues 1 – 450 of hCPEB3. The chemical shifts of the complete IDR of hCPEB3 are reported in the **BMRB (**entry number **50256)**, and the original 2D and 3D spectral data have been deposited in the Mendeley data repository. The assigned 2D ^1^H-^15^N HSQC and 2D ^13^CO^15^N spectra of segment 4 are shown in **Sup. Fig. 3** and **Sup. Fig. 4**, respectively. The 2D ^1^H-^15^N HSQC spectra of segments 1, 3, 4, 5, 6, 7 and 8 are shown in **Sup. Fig. 5**. The similar position of most signals of ^1^H-^15^N in neighboring segments additionally suggests a sparsity of long range interactions under these conditions. Likewise, the majority of the crosspeaks of the same residues in adjacent segments also overlap or are close together in the 2D ^13^CO^15^N spectra of segments 1, 3, 4, 5, 6 and 8 (**Sup. Fig. 6**).

Multiple attempts to express and purify hCPEB3 segment 2, which spans residues 51-150, by recombinant methods were unsuccessful. Nevertheless, all the residues within segment 2 are present and have been characterized structurally in the context of segments 1 and 3. To test if there might be some structure in the neighborhood of residues 91-110 located in the middle of segment 2 and the C- and N-termini of segments 1 and 3, respectively, we studied the conformation of a twenty residue peptide corresponding to this region by NMR spectroscopy. No significant trends towards structure formation were detected (**Sup. Fig. 7**).

The ^13^Cα and ^13^CO conformational chemical shifts (Δδ) of the hCPEB3 segments 1, and 3-8 are plotted in **Sup. Fig. 8**. These data show five segments, comprising residues 1-10, 202-210, 222-234, 238-246 and 346-356, with significantly high Δδ ^13^Cα and Δδ ^13^CO values. Such values are characteristic of partially populated α-helices and are examined in detail in the following paragraphs.

### The first residues of hCPEB3 adopt a partly populated *α*-helix which precedes the Q-rich stretch

The conformational chemical shifts point to the formation of partly populated (20 %) α-helix in the first ten residues of the protein (**Figure 2 A,B**). Increased conformational chemical shifts are observed at 5 °C, reflecting a higher amount of helical structure upon cooling. Standard ^1^H-detected ^15^N relaxation measurements detect that these residues are the most rigid part of segment 1 (**Figure 2 C,D**). These results are corroborated by ^13^C-detected ^15^N relaxation experiments (**Sup. Fig. 9**).

**Figure 2:**
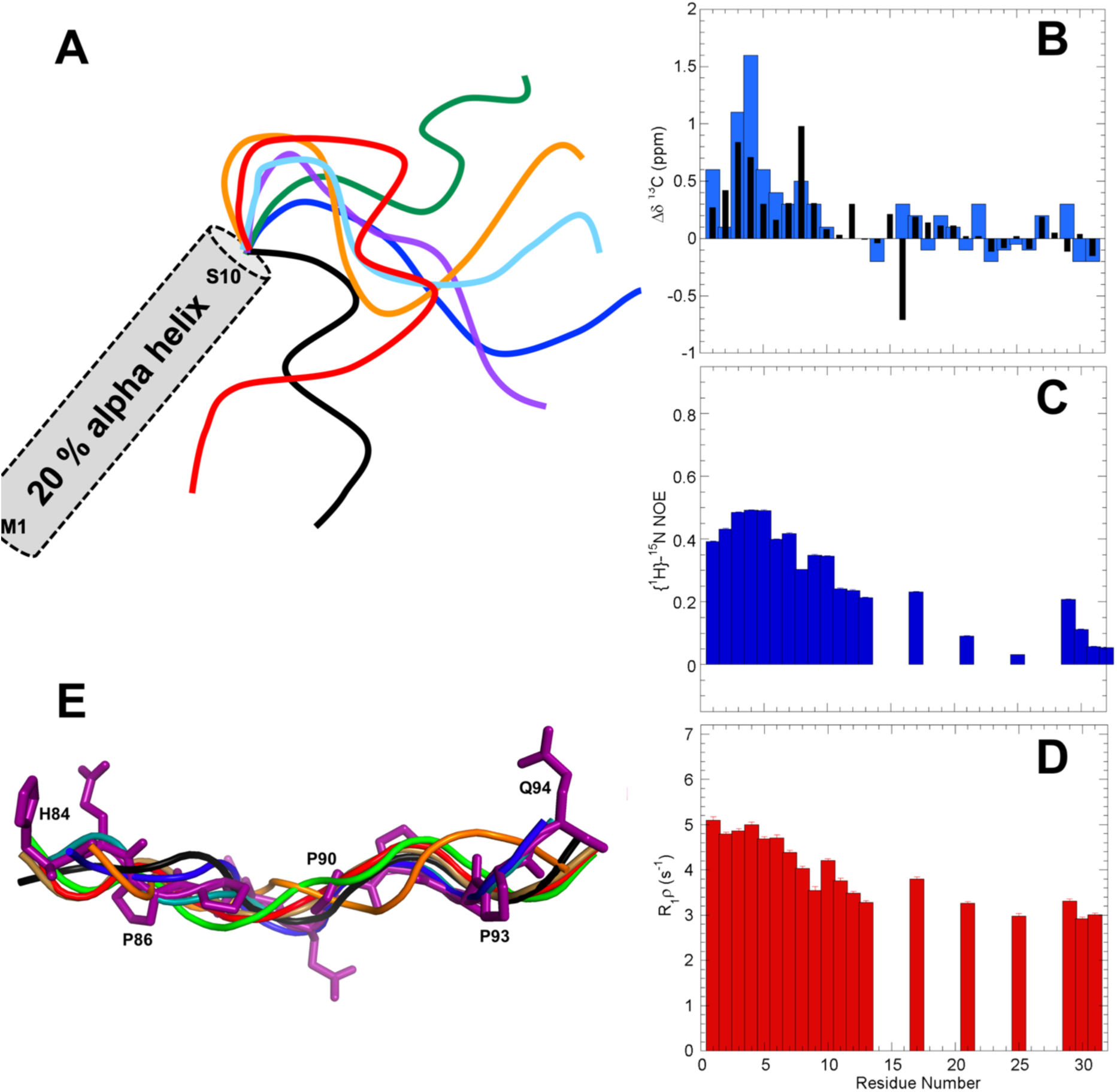
The N-terminal 25 residues of hCPEB3 adopt a hydrophobic α-helix followed by a disordered polyQ segment flanked by PQP mini-breaker motifs. The N-terminus of hCPEB3 contain an α-helix-forming and a disordered amyloidogeneic Q_4_RQ_4_ segments, which are separated by PQP mini breaker motifs. (A) Schematic representation as a gray cylinder of partial (20%) α-helix formation by the first ten residues of hCPEB3. The disordered conformational ensemble of residues 11 -32 is represented curved lines colored purple, blue, cyan, green, orange, red and black . (**B**) ^13^Cα (**blue**) and ^13^CO (**black**) conformational chemical shifts indicate a 20% population of helix at 25 °C. Uncertaintes in the conformational chemical shifts (Δδ) are 0.02 and 0.10 ppm for ^13^CO and ^13^Cα, respectively. (**C**) {^1^H}-^15^N NOE and (**D**) R_1_ρ relaxation measurements indicate that this helical conformation is less mobile than the polyQ segment at ns/ps and μs/ms timescales, respectively at 25 °C. Error bars are shown in panels (**C**) and (**D**) but are small as the estimated uncertainties are <0.01 for the hNOE and < 0.1 s^-1^ for R_1_ρ. Missing values in panels **C** and **D** are due to overlap of ^1^H^15^N peaks or a lack of ^1^H^15^N signals in the case of proline residues. (**E**) Eight representative backbone conformers, colored purple, blue, teal, green, amber, orange, red and black, of the proline rich segment, H84-Q94, featuring a PPII helix that spans residues P86-Q94. All heavy atoms are shown for the purple conformer.

The α-helix detected for these first residues extends N-terminally into the His/Tev tag. To rule out a possible structure-promoting effect on segment 1, we tried to remove it by proteolytic cleavage with the TEV protease. Multiple attempts failed, which suggests that the helix spanning the last residues of His/Tev tag and the first residues of the hCPEB3 IDR is present and impedes the proteolytic cleavage. Therefore, we characterized a dodecamer peptide whose sequence corresponds to the first twelve residues: M_1_QDDLLMDKSKT_12_, of the hCPEB3 IDR. The observation of a series of weak ^1^HN_i_ - ^1^HN_i+1_ nuclear Overhauser enhancement (NOE) crosspeaks reveals that this peptide has a slight tendency to form α-helix in aqueous buffer (**Sup. Fig. 10A**). Fluorinated alcohols like trifluoroethanol (TFE) and hexafluoroisopropanol (HFIP) are known to increase the population of helical conformations in peptides which have an α-helix forming tendency, but not in peptides which prefer to adopt β-strands or random coil [29]. In the presence of 20% HFIP, the population of helix in this peptide increases strongly, based on the observation of stronger and more numerous NOE crosspeaks as well as ^1^Hα and ^13^Cα conformational chemical shifts (**Sup. Fig. 10B**). These findings evince that the first 12 residues of hCPEB3 do tend to adopt an α-helix.

Interestingly enough, the polyQ segment, Q_16_QQQRQQQQ_24_, does not form an α-helix or a β-strand and appears to be thoroughly disordered and flexible (**Figure 2A,B**). A construct spanning residues 1-200 of hCPEB3, which contains the Q_4_RQ_4_ motif, plays a role in hCPEB3 amyloid formation as indirectly evidenced by the anti-amyloid action of the polyglutamine binding peptide 1 (QBP1) [22]. This polyQ segment is preceded and followed by Pro-Gln-Pro residue triplets (P_13_QP_15_ and P_25_QP_27_). Considering the inhibitory effect of proline residues previously observed for polyQ amyloid formation in Huntingtin by Wetzel and co-workers [30], it is likely that these PQP mini-motifs check amyloidogenesis by the polyQ segment. The first 100 residues also contain a predicted SUMOylation site [22] at Lys 47 and ends with a proline-rich segment P_86_PQQPPPPQEPAAPG_100_, which is associated with solubility. Whereas recently reported NMR criteria [31], allow us to rule out that this stretch folds into a stable polyproline II (PPII) helical bundle, the steric limitations of polyproline segments mean that residues 86-93 adopt an isolated, partly populated PPII helix (**Figure 2E**). In fact, the consecutive proline residues show a distinct pattern of conformational chemical shifts; namely +0.6 ppm, -1.0 and -0.3 for ^13^Cα, ^13^Cβ and ^13^CO, respectively (**Table 3**, **Sup. Fig. 11**). Not observed in isolated proline residues, we advance that they are hallmarks of a PPII helical conformation.

### Residues 101 - 200 of hCPEB3 contains a rigid nonpolar segment and a PPII helix

Regarding residues 101 – 200, no strong trends to adopt α-helical or β-structures are detected. Nevertheless, the stretch composed of residues, W_111_STGTTNAVEDSFFQGITPVNGTMLFQNF_139_ which contain numerous aliphatic and aromatic residues, shows relatively high rigidity, both on fast ns/ps as well as slower μs/ms timescales (**Sup. Fig. 12)**. This finding is interesting considering that this relatively hydrophobic segment also appears to be essential for hCPEB3 amyloid formation *in vitro* [22] and very recently it has been reported to form amyloid in mouse CPEB3 [32]. In addition, the stretch of residues 161-190: Q_161_HHQQPPPPA_170_PAPQPAQPAQ_180_PPQAQPPQQR_190_, has a very high Q/P content and Pro and Gln are the residues with the highest intrinsic tendencies to adopt PPII helices [33]. The consecutive proline residues, P_166_PPPAPAPQP_175_, also display the characteristic PPII pattern of conformational chemical shifts (**Sup. Fig. 11 ABC**) seen for residues 86-93. Although more weakly than long stretches of pure polyproline [34][35], a synthetic peptide corresponding to residues P166-P175 of hCPEB3 binds to human Profilin 1, a known mediator of interactions with actin (**Sup. Fig. 11 D**).

### Residues 201-300 contain three *α*-helical segments and a disordered (VG)_5_ segment

Significant ^13^Cα and ^13^CO chemical shift deviations with respect to values predicted for a statistical coil, for three residue segments spanning **<u>A_202_QRSAAAY</u>_21_**_0_ GHQPIMTSKP_220-_**<u>S</u>_221_<u>SSSAVAAAA_230_AAAAA</u>**) SS**<u>ASS_240_SWNTHQ</u>** SVHAA_250_ (**Figure 3A**). These results indicate that the three segments of underlined residues adopt partially populated α-helices. Based on the magnitude of the conformational chemical shifts, the helical populations are different; being about 30% for the A_202_-Y_210_ α-helix, 80% for the S_222_-A_234_ α-helix and 20% for the A_238_-Q_246_ α-helix at 5 °C; whereas these populations decrease at 25 °C to approximately 10%, 40% and 15%, respectively, they are still significant (**Figure 3B**). The presence of the first and second helices are confirmed by ^1^HN-^1^Hα coupling constants (**Sup. Fig. 13**). Moreover, analysis with TALOS+, which predicts secondary structure taking into account ^13^Cβ, ^15^N and ^1^Hα chemical shifts in addition to ^13^Cα and ^13^CO, confirms the presence of these three helical segments and structural calculations with CYANA suggest that the three helices do not tend to adopt a preferred alignment relative to each other (data not shown). The helices are not especially rigid on fast ps-ns time scales (**Figure 3C**) or the slower μs-ms time domain (**Figure 3D**) at 25°C, but do show a heightened stiffness at 5 °C (**Sup. Fig. 12**). Helical wheel projections (**Sup. Fig. 14**) suggest that different interactions contribute stability to these α-helices. Gly 211 and His 212 are positioned to stabilize the A_202_-Y_210_ α-helix by a C-capping motif [36]. Whereas Ala has a very high intrinsic helix forming propensity, the propensity of Ser is low [37]. In this segment, however, the Ser residues are positioned at the N-terminus of the α-helices, where adding negative charge via phosphorylation would increase the helical population, considering the well-known stabilizing effects of charge / macrodipole interactions and N-capping H-bonds [38]. Although this segment’s insolubility thwarts attempts to test directly the impact of phosphorylation using synthetic peptides, we note that these Ser residues are placed at the positions where phosphorylation is expected to increase α-helix stability the most [39]. Moreover, at neutral pH, where phosphoserine carries two negative charges, the stabilization is substantially greater than at pH 4, where it carries one [39]. The last α-helix, A_238_-Q_246_, is less populated, but its stability might increasee if W242 were to engage in long-range interactions, such as with the hydrophobic or cationic residues of the first α-helix, *i.e.* <u>M</u>QDD<u>LLM</u>D<u>K</u>S<u>K</u>T. To test this possibility, we studied two polypeptides containing the M_1_-T_12_ and A_238_-Q_246_ helical segments with and without an N-terminal Dansyl group, connected by a flexible (Gly)_4_ linker. The results of FRET and 2D NMR spectroscopy evince that this polypeptide adopts a conformational ensemble significantly more compact than a statistical coil, but that the helix contents are not significantly altered (**Sup. Fig. 15).**

**Figure 3.**
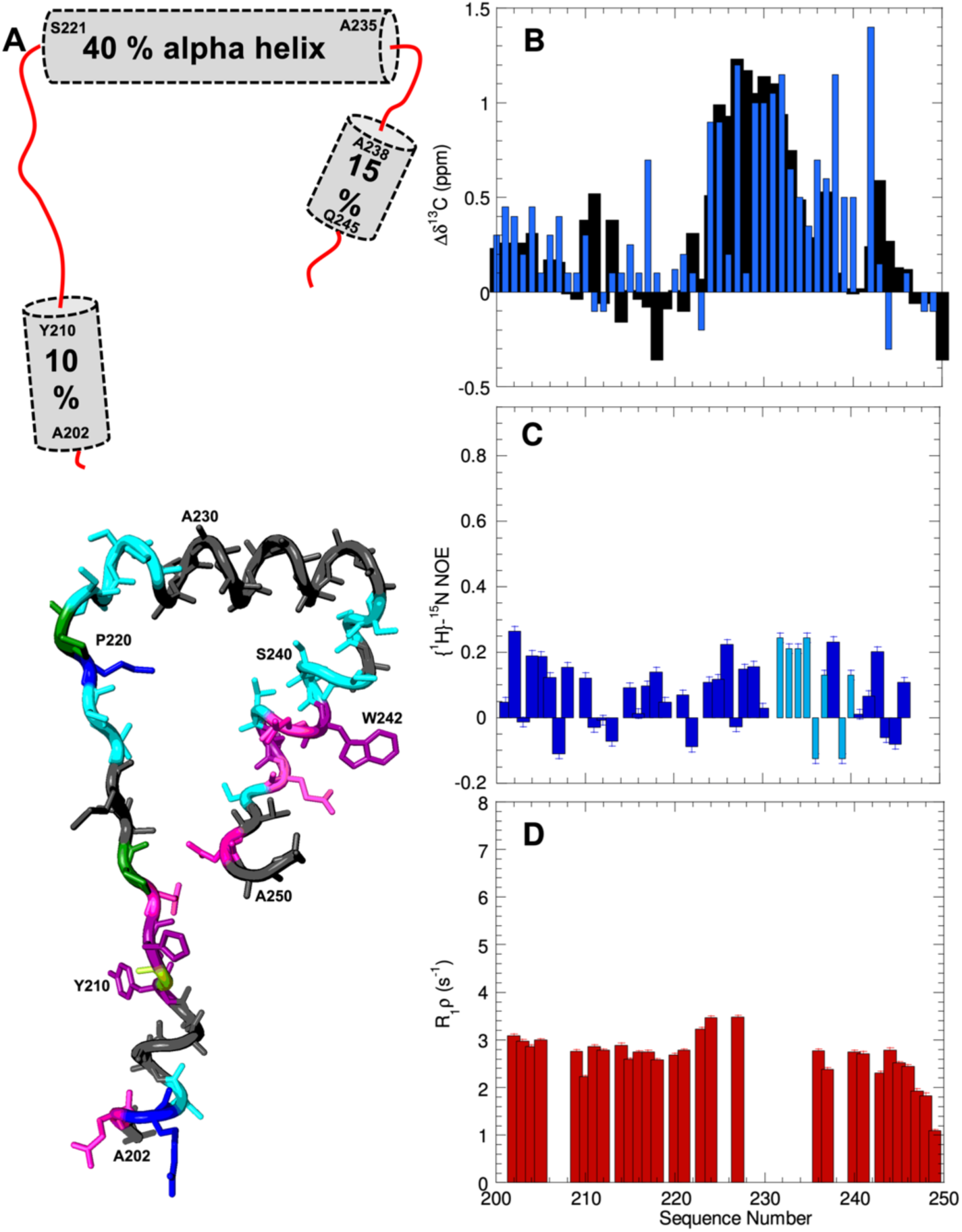
Residues 201 – 250 adopt three partial populated α-helices. **A.** (*Top*) Schematic representation as gray cylinders of the three parcially populated helices present in residues 200 - 250. (*Bottom*) One conformer with all three α-helices is shown; residues are colored: cationic residues (R & K) = **blue**, aromatics (F, Y, W & H) = **purple**, anionic (E & D) = **red**, aliphatic (A, I, L, M) = **dark gray**, amyloidogenic (N & Q) = **magenta**, hydroxyl bearing (S & T) = **cyan**, and proline = **green**. **B.** ^13^CO (**black**) and ^13^Cα (**blue**) conformational chemical shifts (Δδ) of residues 201-250 at 25°C. Note that the second α-helix which contains nine consecutive Ala residues has a relatively high helical population. Uncertainites in the conformational chemical shifts (Δδ) are 0.02 and 0.10 ppm for ^13^CO and ^13^Cα, respectively. **C.** {^1^H}-^15^N NOE ratios. Values shown in dark blue are of individual ^1^H^15^N resonances; those in light blue correspond to overlapped peaks. **D.** R_1ρ_ values reveal the ps/ns and ms/ms time scales. Significantly higher {^1^H}-^15^N NOE ratios and R_1ρ_ values are observed for these residues at 5 °C (**Sup. Fig. 10**).

One of the most striking features in hCPEB3’s sequence is a short dipeptide protein motif (Val-Gly)_5_ spanning residues 271 – 281, which is reminiscent of longer (Ala-Gly)_N_ and (Pro-Gly)_N_ and (Arg-Gly)_N_ dipeptide repeat proteins encoded by mutant C9orf72 which have been implicated in ALS [40]; [41]. Our *in silico* analysis identified this segment as having a high potential to form amyloid [22]. In the context of hCPEB3, however, this segment is among the most disordered and flexible of all the zones of the IDR (**Sup. Fig. 8, 12**). Just beyond the (VG)_5_ segment, there is a stretch of 15 residues: S_284_PLNPISPLKKPFSS_298_, whose NMR parameters indicate disorder and flexibility (**Sup. Fig. 8 & 12**). Nevertheless, this stretch contains four Ser residues reported to phosphorylated by protein kinase A (PKA) or calcium/calmodulin-dependent protein kinase II [42] (**Table 1**), and therefore might be important for the transition between short- and long-term memory. Residues P_303_-PKFP**R**AAP_311_ are proline rich. Predictions suggest that Arg 308 can be methylated (**Table 1**). This modification, whose impact has not been probed here, was reported to fortify cation–π interactions, reduce interactions with RNA and destabilize condensates in other proteins [43].

### The residues forming the Nuclear Export Signal (NES) show a marked tendency to adopt *α*-helical structures

Significant conformational chemical shifts were also observed for residues L349-L353 which form the NES (**Figure 4**) indicating the presence of helical structure. Using the ^13^CO, ^15^N, ^1^HN, ^13^Cα and ^13^Cβ chemical shift data as input, a family of conformers was calculated using the programs TALOS+ and CYANA for residues P333-P363. This 31-residue segment is rich in aromatic (five) and aliphatic (six) residues, which is unusual for a disordered polypeptide. The resulting structures reveal that residues L346-L349 adopt one turn of α-helix and residues S352-M356 form a short α-helix (**Figure 4A**). It is notable that this conformer positions five nonpolar residues: L346, L349, L353, M354 and I357 on the same face of the α-helices. Y341, the putative phosphorylation site, is in an extended portion of the backbone and would be accessible for this post-translational modification (PTM). Whereas the conformational ensemble will contain many other structures, based on conformational chemical shifts as illustrated by the Δδ^13^Cα & Δδ^13^CO values shown in **Figure 4B** the α-helical population is about one third. The presence of rigid conformers is corroborated by relatively high {^1^H}-^15^N NOE ratios (**Figure 4C**) and elevated transverse relaxation ratios (**Figure 4D**).

**Figure 4.**
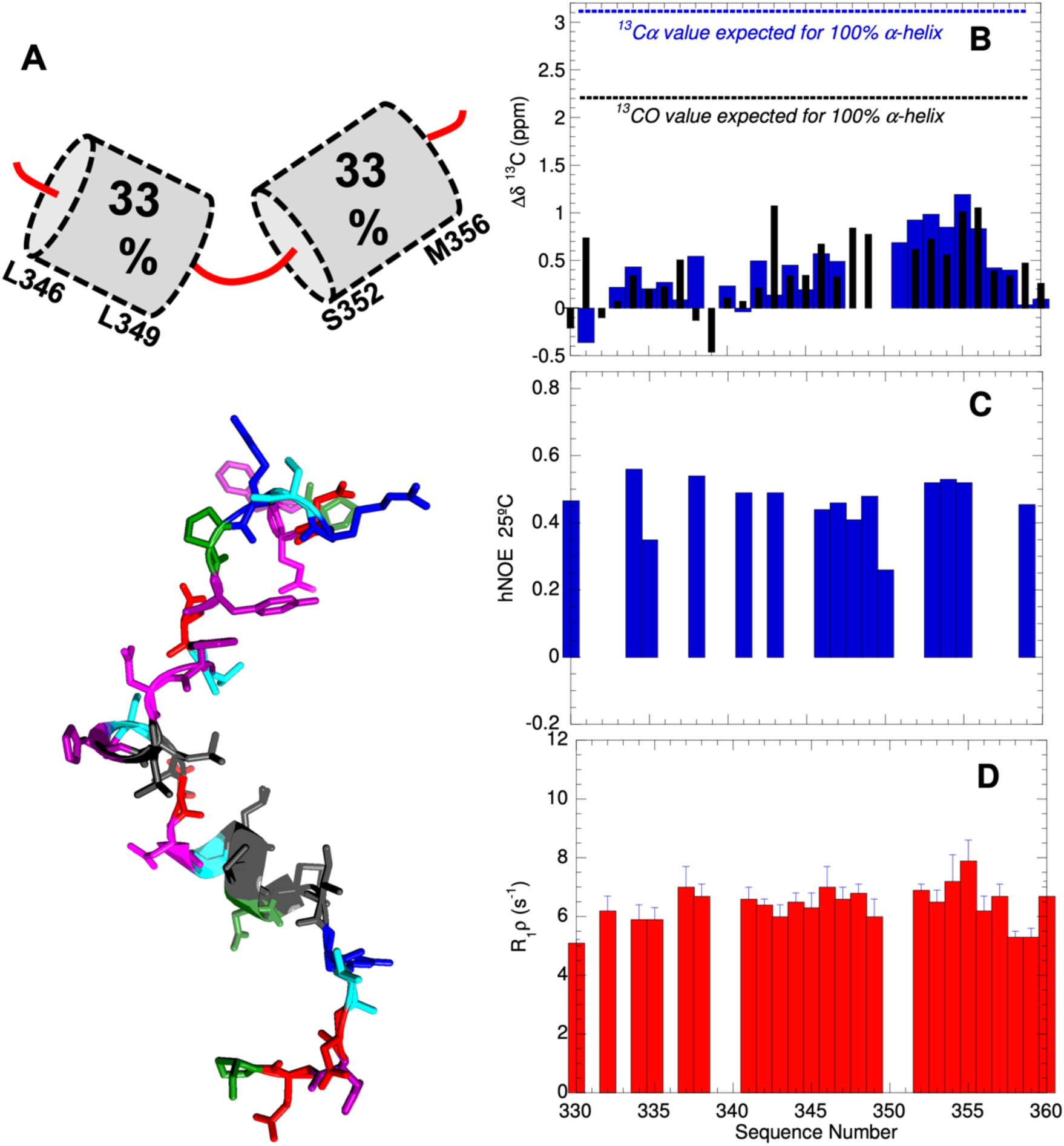
Conformation of the NES and nearby phospho-Tyr site. **A**. (*Top*) Residues L346-L349 and S352-M356 adopt two short, partially populated α-helices. (*Bottom*). Representative conformer of shown with cationic residues (R & K) colored **blue**, aromatics (F, Y & H) = **purple**, anionic (E & D) = **red**, aliphatic (I, L, M) = **dark gray**, amyloidogenic (N & Q) = **magenta**, hydroxyl bearing (S & T) = **cyan**, and proline = **green**. Spiral ribbons mark the helical segments spanning residues 346 – 349 and 352 - 356. **B**. Conformational chemical shifts of ^13^Cα (**blue** bars) and ^13^CO (**black** narrow bars) afford detection of helical conformations. Uncertaintes in the conformational chemical shifts (Δδ) are 0.02 and 0.10 ppm for ^13^CO and ^13^Cα, respectively. **C.** {^1^H}-^15^N NOE ratios of 0.85 and -0.20 are indicative of high rigidity and flexibility, respectively, on ps-ns time scales. **D**. Higher R_1ρ_ rates are diagnostic of rigidity on μs-ms time scales.

Beyond the NES α-helix, no segments with preferred secondary structure are detected. The last residues of the segment 8 construct: S_426_-RKVFVGGLPPDIDEDEITASFRRF_450_, belong to the RRM1 domain. According to the 3D structure [18] residues K428-G432 adopt a β-strand and residues E440-R449 form an α-helix in the context of the complete RRM1 domain. Here these segments appear to be largely disordered. After a proline-rich zone ending around residue 380, the next fifty residues have a higher content of nonpolar residues and tend to be more rigid (**Sup. Fig 12**). Residues 400 – 412: SHGDQALSSGLSS contain five Ser residues reported to be phosphorylated [42], **Table 1**). Like the four Ser residues of the 284-291 stretch mentioned above, this could be an element involved in the structural basis of memory consolidation.

## Discussion

Beyond the fundamental biological interest in comprehending the molecular basis of long-term human memory, the atomic-level understanding of the structural transitions involved in CPEB aggregation could also guide the development of treatments for Post-Traumatic Stress Disorder and other diseases associated with the storage of “pathological memories” [44]. Furthermore, like its homologs in *Aplysia* and *Drosophila*, hCPEB3 resembles the abundant and diverse superfamily of RNA-binding proteins that contain RRM and/or ZnF domains as well as intrinsically disordered prion-like regions, such as Fused in Sarcoma (FUS) or Transactive Response DNA Binding Protein of 43 kDa (TDP-43). FUS and TDP-43 are essential proteins, but their anomalous aggregation has been implicated in amyotrophic lateral sclerosis (ALS) and frontotemporal dementia (FTD) [45]. In addition, TDP-43 has been linked to the recently described Limbic-Predominant Age-Related TDP-43 Encephalopathy (LATE) [46]. Thus, the comparison of the physiologically relevant amyloid formation, and the dynamics at the monomer level, by hCPEB3 versus the aberrant amyloid formation by TDP-43 may reveal why the latter can become pathological.

The biophysical analysis of the full length IDR of hCPEB3 shows that it lacks stable secondary structure. Compared to well-ordered protein domains, disordered regions tend to change their amino acid sequence much more rapidly over the course of evolution due to a lack of structural constraints for folding [47]. In hCPEB3, the N-terminal α-helix and the 350’s (NES) α-helix as well as its accompanying putative phosphoTyr site, are well conserved throughout vertebrate CPEB3’s, which suggests that they are important for biological function (**Sup. Fig. 2**). Additional segments, like the stretch of hydrophobic residues key for amyloid formation [22] and a cluster of three Trp residues (W242, W252 and W259) are also conserved from mammals to fish. In contrast, the Gln-rich segment at the N-terminus, the Pro-rich ’breaker’ regions and the Ala-rich helices are well conserved in mammals but not across all vertebrates. In some lower vertebrates, there is an alternative Q-rich region positioned after the 100’s hydrophobic segment. The relatively rapid evolution of these elements could be related to the development of the mammalian brain as compared to those of lower vertebrates. By contrast, the ability to move the poly-Q segment or substitute it for hydrophobic amyloidogenic segment highlights the cassette or modular nature of PLDs, which was previously established for the *Drosophila* CPEB homolog [5].

Over the past decade, the PLD of several proteins involved in transcription regulation and/or RNA metabolism have been characterized. Some of these, including Huntingtin [48], the Androgen Receptor [49], Ataxin-7 [50] and TDP-43 [51] (**Table 2, Sup. Fig. 14**) have a hydrophobic, helix-forming segment which precedes the Q- or Q/N-rich amyloid-forming segment. These hydrophobic helices have been proposed to promote intermolecular associations, either through the formation of coiled-coils [52] [53] or by forming biomolecular condensates via liquid-liquid phase separation [54]. In the case of Huntingtin, the poly-Q segment is followed by a stretch of poly-P, which will be locked in a PPII helical conformation. This poly-P stretch has been shown to strongly slow Huntingtin amyloid formation *in vitro* [30]. However, poly-Q expansions in Huntingtin protein [48] and the Androgen Receptor [55] are known for their ability to form toxic aggregates in Huntington’s disease and in spinobulbar muscular atrophy, respectively. Interestingly enough, Huntingtin homolog in *Aplysia* has been proposed to play a role in memory consolidation [56].

**Table 2:**
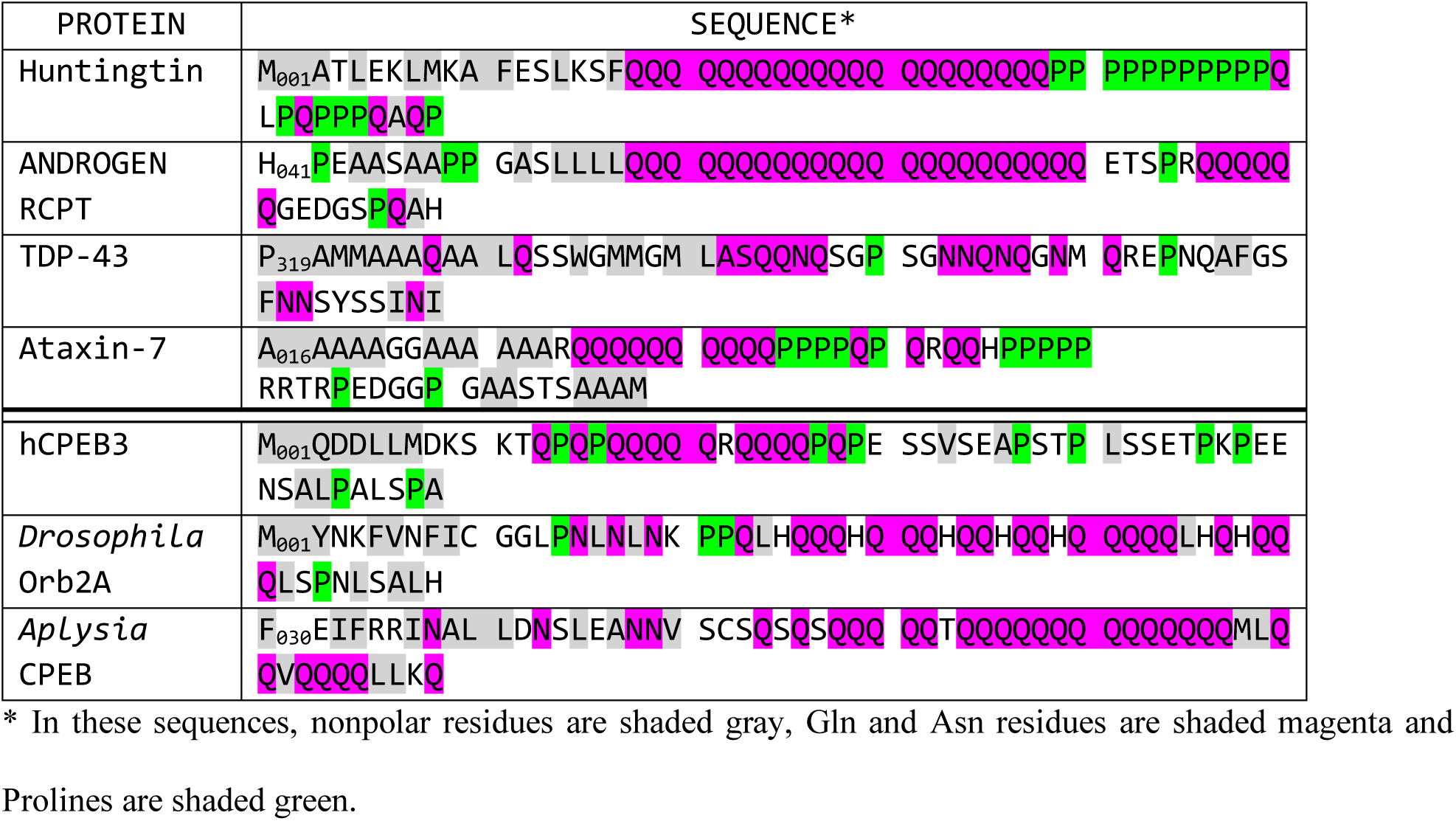
Hydrophobic helices and Pro rich stretches modulate amyloid formation by Q-, Q/N-rich segments.

### PPII helices may regulate amyloid formation and mediate interactions with actin

Whereas hCPEB3 and its homologs Orb2 and *Ap*CPEB have α-helix destabilizing residues between the N-terminal helix and the amyloidogenic Q-rich segment, such as proline in hCPEB3 and Orb2 or serine and valine in the case of *Ap*CPEB, no such α-helix busters are present in the analogous segments of Huntingtin, the Androgen Receptor or TDP-43 (**Table 2**). This allows the poly-Q segment to interact with the α-helix through the formation of sidechain to main chain hydrogen bonds in the case of the Androgen Receptor [55] and to form part of the amyloid structure as proposed recently for TDP-43 by Eisenberg and collaborators [57] and present in ALS/FTLD patient brains [58]. Based on these observations, we suggest that the presence of α-helix breakers between the α-helix and the poly-Q(/N) segments in functional amyloids and their absence in pathological amyloids is a fundamental difference that could be a molecular basis of their radically different toxicities.

PPII helices, well known in collagen and certain glycine-rich proteins [59], also play key roles in mediating protein-protein interactions in biomolecular condensates [60]. Whereas conformation chemical shifts have proven to be extremely useful tools to identify α-helices and β-strands in folded and disordered proteins, conformational chemical shifts for PPII helices were less known. The first 450 residues of hCPEB3 include no less than 66 proline residues (15% of the total). We hypothesize that segments such as G_166_PPPPAPAPQP_175_, may bind profilin, which has specific domains to interact with proline-rich polypeptides [34] and to mediate interactions with the actin, whose levels rise at the synapse following CPEB3 aggregation [13]. On the basis of the thorough set of chemical shifts obtained here, including Pro ^15^N assignments which are rare in the literature, we propose a pattern of conformational chemical shifts that define PPII helices (**Table 3**). Combined with the recently published standards for glycine-rich PPII helical bundles [31], these values should aid the detection of PPII helices in biomolecular condensates and the elucidation of their roles in physiology and pathology.

**Table 3:** Conformational Chemical Shifts for α-helices, β-strands and PPII helices.

| | α-helix | β-strand | Gly-rich<br>PPII helical bundle | Isolated Pro-rich<br>PPII helix |
| --- | --- | --- | --- | --- |
| $\delta\Delta\ ^{13}\text{C}\alpha$ | 3.1<sup>a</sup> / 2.9<sup>b</sup> | -1.5<sup>a</sup> / -1.8<sup>b</sup> | -0.6<sup>d</sup> | +0.6<sup>e</sup> |
| $\delta\Delta\ ^{13}\text{C}\beta$ | -0.4<sup>a</sup> / -0.8<sup>b</sup> | +2.2<sup>a</sup> / +2.7<sup>b</sup> | +0.3<sup>d</sup> | -1.0<sup>e</sup> |
| $\delta\Delta\ ^{13}\text{CO}$ | 2.2<sup>c</sup> / 2.2<sup>b</sup> | -2.2<sup>c</sup> / -2.1<sup>b</sup> | -0.2<sup>d</sup> | -0.3<sup>e</sup> |

**a.** from Spera & Bax \(1991\) [100].
**b.** from Wang and Jardetzky \(2002\) [101] for alanine.
**c.** calculated from Wishart & Skyes \(1994\) [102].
**d.** from Treviño \*et al.\* \(2018\) [31].
**e.** This study.

### STAT5B binding may occlude hCPEB’s NES leading to nuclear retention

The calculated conformers for the NES showed the preferred formation of α-helices which orient five hydrophobic residues on the same side (**Fig. 4**); this is a common feature in nuclear export signals [61]. CPEB3 constantly shuttles in and out of the nucleus and its subcellular localization has important consequences as it regulates gene transcription via association with STAT5B in the nuclei of neurons as well as mRNA translation in dendritic spines. Neural stimulation leads to hCPEB3, which is normally predominately cytoplasmic, to be concentrated in the nucleus [17]. That earlier study also revealed that the region spanning residues 639 – 700 of STAT5B binds the N-terminal IDR of hCPEB3 [17]. To learn about the nature of this interaction, we examined the structure of STAT5B (PDB: 6MBW) [62]. Remarkably, this structure reveals that the key hCPEB3-interacting residues in STAT5B form an SH2 domain. Based on this, we advance that hCPEB3 may contain a phospho-Tyr residue that is recognized and bound tightly by STAT5B. Considering the results of previously reported motif analyses [63][64], the most plausible site for phospho-Tyr modification in hCPEB3 is Tyr341, as this residue is bordered by SRPY_341_DTF which closely matches a consensus substrate “[E/V/D]-[G]-[I/P/L]-Y-E-X-[F/T/V]” of Scr, which is one of the most relevant tyrosine kinases. Based on this proposed PTM site, and our finding that the preferred conformation of nearby NES includes an α-helix spanning residues L349-L353 *(vide supra*), we can advance a speculative working hypothesis to account for how STAT5B binding retains hCPEB3 in the nucleus; namely binding of phospho-Tyr341 by the SH2 domain of STAT5B may occlude the neighboring NES of hCPEB3. This might explain why hCPEB3 becomes concentrated in the nucleus under conditions of neuronal stimulation [15]. Recent studies have shown that certain mutations in the nuclear localization and export signals or defects in nucleocytoplasmic transport of TDP-43, another aggregation-prone, RNA-binding protein, lead to an aberrant aggregation [65]. Although beyond the scope of this study, future research could test whether mutations in the NES of hCPEB3 affect transport and lead to long-term memory defects.

### Towards an atomic-level description of the first steps in hCPEB3 IDR self-association

We suggest a speculative working hypothesis for hCPEB3 conformational changes during memory consolidation (**Figure 5**). Initially, the first 200 residues of hCPEB3, which are necessary and sufficient for aggregation and amyloid formation *in vitro* [12][22], are mostly disordered except for the relatively stable α-helices formed by residues 222-234 and modestly populated α-helices at the N-terminus and spanning residues 202-212 and 237-245 (**Figure 5A**). The Q_4_RQ_4_ motif and hydrophobic segment **F_123_FQGIT**-PVNGT-**MLFQNF_139_** are initially disordered and premature amyloidogenesis is discouraged by SUMOlyation [14] and proline breaker motifs.

**Figure 5.**
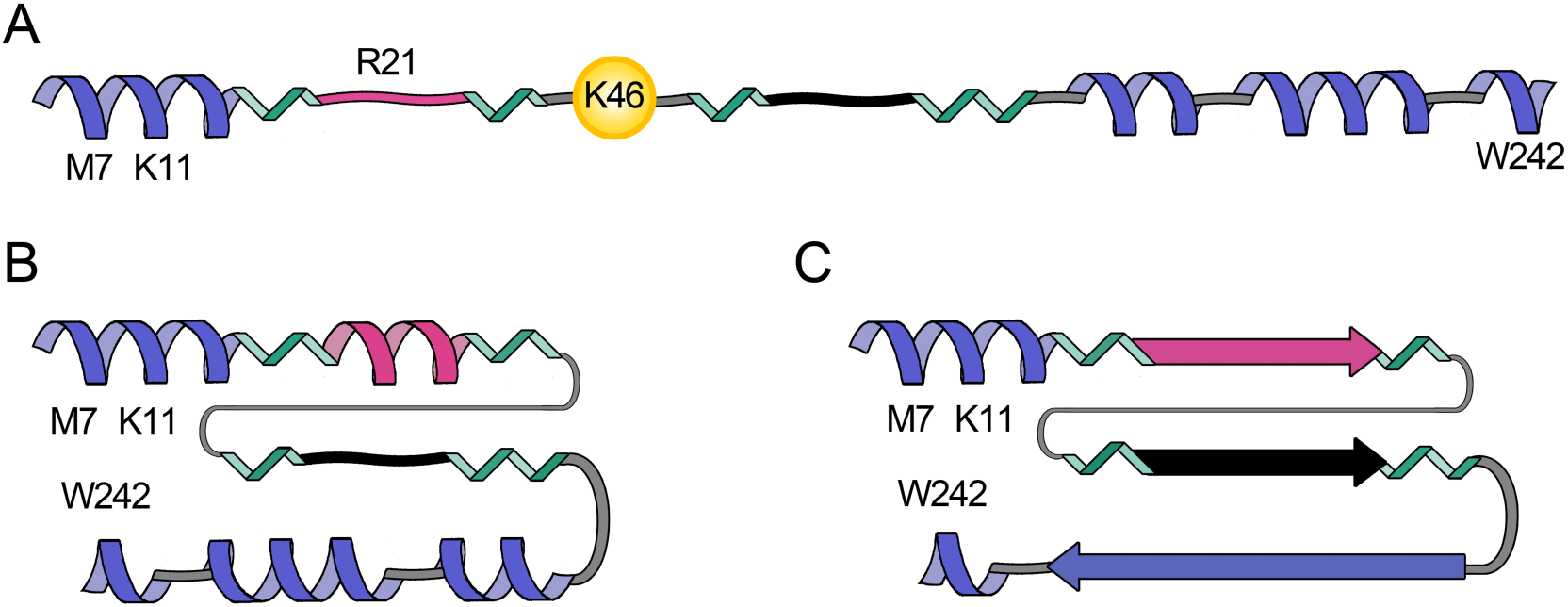
Working hypothesis for hCPEB3 structural changes during memory consolidation. **A.** The first 250 residues of hCPEB3 contain 4 α-helices (blue spirals), the first and third α-helices are relatively stable. Proline-rich segments (green) and SUMOylation as putatively to occur at K46 by *in silico* methods (Ramírez de Mingo *et al.,* 2020) prevent premature association and amyloid formation by the Q_4_RQ_4_ segment (magenta) and the hydrophobic motif (**black** squiggle). **B.** Following deSUMOlyation and putatively phosphorylation, association between the fourth and first helices could occur, strengthened by hydrophobic and cation-π interactions. The Q_4_RQ_4_ segment may adopt an α-helix and associate with the Ala rich α-helices to form a coiled-coil. The structural transformations may well enhance intermolecular contacts within the dendritic P-body like granule, leading to gelification and eventually, amyloid formation. **C.** The final amyloid could be composed of the Q_4_RQ_4_ segment and hydrophobic tract, and possibly the Ala-rich segments. The final configuration of the polyPro segments (**green**) may promote profilin binding and the initiation of a more robust actin filament network.

Following deSUMOlyation and other putative PTMs such as phosphorylation, association among the α-helices into more compact ensembles, as suggested by FRET results (**Sup. Fig. 15**), would become possible, and might be strengthened by hydrophobic interactions between Met 7 and Trp 242 or cation-π interactions between Lys 11 and Trp 242 (**Figure 5B**), analogous to the long range contacts seen in a TDP-43 amyloid [66]. The increased production of hCPEB3 upon neuronal stimulation [12] as well as the proximity of the serine and alanine rich α-helix (residues 222-234) may promote α-helix formation by the QQQQRQQQQ motif. These could then associate to form a coiled-coil, as has been demonstrated in model polypeptides [67] [52] [68]. Moreover, completely helical polyalanine peptides, some of which come from proteins linked to polyalanine expansion diseases, were reported to promote coiled-coil mediated aggregation although in some case no conversion into β-sheet structures was observed [52] [69]. Although future studies are necessary to provide additional evidence and test plausible hybrid aggregation mechanisms with several elements of structure acting in parallel, this model is already supported by analogous results on *Ap*CPEB [4] [53] and polyalanine expansions [69], which led to the proposal of similar mechanisms.

These events would reinforce intermolecular contacts within the dendritic P-body like granule [20] leading to gelification and, eventually, amyloid formation (**Figure 5C**). The final amyloid structure could be comprised of the Q_4_RQ_4_ motif and the hydrophobic segment as evidenced by hCPEB3 fibril formation kinetics. A construct spanning residues 1-200 of hCPEB3, which contains the Q_4_RQ_4_ motif, plays a role in hCPEB3 amyloid formation as indirectly evidenced by the anti-amyloid action of the polyglutamine binding peptide 1 (QBP1) [22] . In the case of the hydrophobic segment, its role in mouse CPEB3 amyloidogenesis has recently been thoroughly corroborated [32]. In addition, the poly-A segment of helices 202-212 and 222-234 may also transform into amyloid (**Figure 5C**). Such α-helix to amyloid conformational transformations have been previously described in alanine-rich polypeptides as diverse as a fish antifreeze protein [70] and a synthetic (Ala)_10_-(His)_6_ hexadecapeptide [71], and could form pathological amyloids in a number of reported diseases involving expansion of poly-A tracts [72].

Residues 217-284 of CPEB3 have been reported to be key for interactions with actin [13]. Considering this, the hypothetical associations among the α-helices formed by residues M_1_-T_12_ and residues A_202_-Q_246_ suggested in the preceding paragraphs could dispose the PPII helices formed by residues P_86_-Q_94_ and P_166_-P_175_ (**Sup. Fig. 10**) to bind profilin. This protein contains a second binding site specific for actin and promotes the formation of actin filament networks [73]. Such actin networks are known to become more extensive and robust as a dendritic bud strengthens during memory consolidation [74].

## Materials and Methods

### 1. Materials

^15^NH_4_Cl and ^13^C-glucose were purchased Tracertec (Madrid, Spain), D_2_O, was a product of Euroisotop, deuterated acetic acid was from Sigma/Aldrich and 4,4-dimethyl-4-silapentane-1-sulfonic acid (DSS) as the internal chemical shift reference, was from Stolher Isotopes Chemical Company. A twelve residue peptide, called hCPEBpep1, whose sequence corresponds to the protein’s first 12 residues (M_1_QDDLLMDKSKT_12_) and a twenty residue peptide, called hCPEBpep2 with the sequence of residues P_91_PPQEPAAPGASLSPSFGST_110_ in hCPEB3 were purchased from Genscript. hCPEBpep2’s sequence overlaps with the C-terminal of Segment 1 and the N-terminus of Segment 3. A proline-rich peptide acGPPPPAPAPQPam, with acetylated and amidated termini and whose sequence covers residues 166-175 in hCPEB3, two polypeptides with the sequence MQDDLLMDKSKTGGGGASSSWNTHQ (with and without an N-terminal Dansyl group) were also obtained from Genscript. All the peptides were over 95% pure, as assessed by HPLC, and their identities were confirmed by mass spectrometry and NMR spectroscopy. Recombinant Human Profilin 1, produced recombinantly in *E. coli* was obtained from Abcam (reference number ab87760). It is over 95% pure as judged by SDS PAGE.

### 2. Sample Production

Coding mRNAs of hCPEB3 vary in length due to an embedded human delta virus-like ribozyme which slowly splices out introns, leading to the generation of multiple isoforms [75]. It is also noteworthy that Orb2A’s pre-mRNA contains an intron with multiple stop codons which is only spliced out when certain “memorable” stimuli are experienced [9]. Here, the hCPEB3 isoform 2, Uniprot Q8NE35-2 / Genebank CAI14105.1, is studied.

#### Plasmid construction, protein expression and purification

The hCPEB3 IDR, which corresponds to the first 426 residues of the protein whose sequence is shown in **Table 1**, was expressed at eight highly overlapping one-hundred residue segments. To control for end effects as well as to check the reproducibility of the results, each segment overlapped by 50 residues with the preceding and successive segment.

Each segment was cloned into the pET-28a(+) by PCR using the full length human CPEB3-2 in pLL3.7 plasmid as the template kindly provided by Dr. Yi-Shuian Huang [76]. The DNA amplified fragments were digested with XhoI and NheI. Expression of the resulting clones led to fusion proteins containing a His_6_ tag and a TEV NIα protease cleavage site. Thus, each segment studied had the sequence: MGSSHHHHHHSSGLVPRGSHMASENLYFQ, at its N-terminus.

All overlapping segments were expressed in the *E. coli* BL21 Star (DE3) strain using the T7 expression system (Novagene). ^13^C/^15^N isotopic labelling of each segment was done by using a previously published protocol [77]. Briefly, cells were grown in 1 L of LB at 37 °C by shaking at 280 rpm upon reaching optical cell densities at 595 nm (OD_595_) ∼ 0.6-0.7. Cells were pelleted by a 30 min centrifugation at 5000 × g and washed using a M9 salt solution (15.0 g/L KH_2_PO_4_, 34.0 g/L Na_2_HPO_4_ and 2.5 g/L NaCl for 1 L of 5 × M9 salts) excluding nitrogen and carbon sources. Cell pellets were resuspended in 250 mL of isotopically labelled minimal media M9 salt solution supplemented with ^13^C D-glucose 4.0 g/L and ^15^NH_4_Cl 1.0 g/L (Cambridge Isotope Laboratories, Inc.), then incubated to allow the recovery of growth and clearance of unlabeled metabolites. Protein expression was induced after 1 h by addition of IPTG to a concentration of 1 mM. After a 4.5 h incubation period, the cells were harvested.

Cell pellets were lysed with buffer with the following composition: 50 mM NaH_2_PO_4_/Na_2_HPO_4_, 500 mM NaCl, 50 mM imidazole, 6 M guanindium chloride (GdmCl), pH 7.4 and then sonicated. Each recombinant segment was purified by Ni^2+^-affinity chromatography using HisTrap HP purification columns with a FPLC system (ÄKTA Purifier, GE Healthcare) with elution buffer consisting of 50 mM NaH_2_PO_4_/Na_2_HPO_4_, 500 mM NaCl, 500 mM imidazole, 6 M GdmCl, pH 7.4.

If necessary, the pure segments were then incubated with TEV NIα protease O/N at 4°C [78]. The cleaved protein was then subjected to dialysis and recovered. Protein samples were stored at -80°C until use. Eventually, they were desalted to 1 mM DAc, pH 4.0 by gel filtration chromatography using a PD-10 column (GE Healthcare), and concentrated to a final protein concentration of 1.0 – 1.5 mM in 250 μL using a Vivaspin microfiltration device and placed in a 5 mm Shigemi reduced volume NMR tube for measurements. These low pH, low ionic strength conditions have been reported [79] and corroborated [80] to maximize solubility by increasing charge-charge repulsion between protein molecules. α-helix stability depends on pH due to charge-charge interactions, charge-helix macrodipole interactions and intrinsic helix propensities [38] . A consideration of these factors suggests that the effect of increasing the pH from 4 to 7 would be mildly stabilizing for the first α-helix and insignificant for the four remaining α-helices detected in the hCPEB3 PLD.

Non-labeled full length hCPEB3-IDR expression and purification was carried out essentially as described in [22]. Briefly, cells were grown in 1 L of LB medium at 37 °C until reaching an OD_600_ = 0.6 – 0.7 and protein expression was induced for 4 hours by adding IPTG at 1 mM final concentration. Cells were harvested and following sonication, then lysed with buffer 50 mM Na_2_HPO_4_, 500 mM NaCl, 50 mM imidazole, 6 M GdmCl pH 7.4. After centrifugation at 18000 rpm for 45 min, supernatants were purified with Ni^++^ affinity chromatography and elution was performed in buffer containing 50 mM NaPO_4_, 500 mM NaCl, 500 mM imidazole, 3 M GdmCl, pH 7.4. Purified CPEB3-IDD was diluted to PBS, 1 M GdmCl pH 7.4, dialysed against PBS pH 7.4 at 4°C and finally concentrated by using Amicon Ultra-15 centrifugal filters 10 kDa cut-off.

### 3. Sequence alignment

The conservation of vertebrate CPEB3 protein sequences was assessed using the programs T-coffee [81] and Clustal Omega [82] using the default settings. The sequences chosen as representative are human isoform 1 NP_001171608.1, human isoform 2 CAI14105.1, mouse NP_001277755.1, chicken XP_105144323.1, turtle (*Chrysemys picta belli*) XP_005301348.1, frog NP.001015925.1 (*Xenopus tropicalis*) and fish (*Danio rerio*) XP_009305819.1.

### 4. Fluorescence and Circular Dichroism Spectroscopies

Fluorescence spectra on the full length, unlabeled hCPEB3-IDR at ca. 5 µM in 1.0 mM deuterated acetic acid at pH 4.0 were recorded on a Horiba FluorMax 4 instrument equipped with a Peltier temperature control device using a 0.2 sec·nm^-1^ scan speed and three nm excitation and emission slit widths. The excitation wavelength was 280 nm and the emission was scanned over 300 – 400 nm. A series of spectra were recorded at 2, 10, 20, 30, 40, 50, 60 and 70 °C.

To test for binding of a segment of hCPEB3 rich in proline residues to Profilin, fluorescence spectra were recorded at 20 °C on 10 µM human Profilin 1 using two nm excitation and emission slits, a 2 nm·s^-1^ scan speed, an excitation wavelength of 295 nm and emission was recorded from 300 to 400 nm in the absence and presence 4.5 mM of the peptide acGPPPPAPAPQPnm, which corresponds to residues P166-Q175 of hCPEB3, with an N-terminal acetyl and glycine residue and a C-terminal amide group added to avoid end charges. This assay is based on blue shift and enhanced emission of Trp3 and Trp31 of human Profilin 1 as the environment surrounding their indole moieties becomes less solvent exposed upon polyproline ligand binding [34].

Fluorescence spectra on Dansyl-MQDDLLMDKSKTGGGGASSSWNTHQ and the control polypeptide without Dansy, MQDDLLMDKSKTGGGGASSSWNTHQ, were recorded using an excitation wavelength of 295 nm, which is selective for the donor Trp, and scanning the emission over 300 – 580 nm, using a 2 nm·s^-1^ scan speed and slit widths of 2 nm. The final concentration of both polypeptides was matched at 10.0 µM. Spectra were recorded in 100 mM KCl, 20 mM K_2_HPO_4_ / KH_2_PO_4_ (pH 7) with or without the denaturant GdmCl present at a final concentration of 7.4 M. Assuming randomized orientations of the donor and acceptor groups and a Förster distance (*R_0_*) of 23.6 angstroms for the Dansyl / Trp pair, their average distance, <r>, can be estimated using the equation 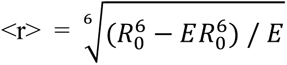, where E is the fraction of energy transferred from donor to acceptor [83].

A Jasco 810 spectropolarimeter fitted with a Peltier temperature control unit was used to record far UV-CD spectra at 5 and 35 °C on the complete hCPEP3 IDD at ca. 5 µM in 1 mM deuterated acetic acid (pH 4.0) using a 1.2 nm bandwidth scanning from 260 to 190 nm in a 0.1 cm quartz cuvette at 50 nm·min^-1^. Eight scans were recorded and averaged for each spectrum.

### 5. NMR Spectroscopy: Instrumentation

All spectra for the hCPEB3 segments were recorded on a Bruker 800 MHz (^1^H) Avance spectrometer fitted with a triple resonance TCI cryoprobe and Z-gradients. The ^1^H chemical shift was referenced to 50 μM DSS measured in the same buffer and at the same temperatures as those used for the CPEB3 segments. Since DSS can sometimes bind to intrinsically disordered proteins) [84], the DSS signal was recorded in an independent reference tube containing the same buffer at the same temperature. The ^13^C and ^15^N chemical shift references values were calculated by multiplying by their respective gyromagnetic ratios with ^1^H; that is Ξ ^13^C / ^1^H = 0.251449530 and Ξ ^15^N / ^1^H = 0.101329118 [85]. NMR spectra were recorded and transformed using TOPSPIN (versions 2.1) (Bruker Biospin).

#### ^13^C-Detection Assignment Strategy

Among the NMR approaches for studying disordered proteins (reviewed by [86], ^13^C-detection has been gaining in popularity as it affords the characterization of proline residues [87][88] and offers superior signal dispersion [89]. Here, to speed and improve the assignment of the backbone, we used a “proton-less” NMR approach for segments 1, 4, 5, 6 and 8 based on 2D CON spectra in which successive ^15^N-^13^CO nuclei correlations are obtained in two 3D spectra called hacacoNcaNCO and hacaCOncaNCO [90]. For segments 7 and 8, which tend to form condensates [22] and seem to be more rigid, this strategy afforded less intense spectra and in particular about 35 CON crosspeaks were missing. Therefore, an additional strategy based on ^13^CO connectivities from 3D HNCO and HNcaCO spectra, as well as ^1^HN and ^15^N connectivities of consecutive residues from 3D HncocaHN and hNcocaNH spectra [91][92] was utilized to check and complete the backbone assignments. The success of this approach seems to be due to a very slow equilibrium between aggregated protein molecules and those remaining in solution, which are detectable by NMR. The latter strategy was also employed for segment 3, which was less soluble. For all segments, further corroboration was obtained by conventional 2D ^1^H-^15^N HSQC and 3D HNCO spectra as well as 3D CCCON to confirm the residue identity and obtain the chemical shift values of ^13^C nuclei of the side chains.

Of the eight segments, only segment 2 failed to yield a soluble sample. Whereas the sequence assignments are complete thanks to the analysis of segments 1 and 3, to test for possible end effects, a 20 residue peptide, hCPEBpep2, corresponding to residues 91-110 of the hCPEB3 sequence was assigned and characterized structurally by 2D ^1^H-^1^H COSY, ^1^H-^1^H TOSCY, ^1^H-^1^H NOESY and 2D ^1^H-^13^C HSQC NMR spectra at 5.0 °C on a Bruker 600 MHz spectrometer fitted with a cryoprobe and Z-gradients. A 2D ^1^H-^15^N HSQC was also recorded on the 800 MHz Bruker spectrometer. Both the ^1^H-^15^N HSQC and the ^1^H-^13^C HSQC spectra were recorded at the natural abundance of ^15^N and ^13^C. The program NMRFAM-Sparky [93] was used to facilitate manual spectral assignment. The NMR spectral parameters are summarized in **Sup. Table 1**.

Theoretical chemical shift values (δ_coil_) for statistical coil ensembles were calculated using the parameters tabulated by [94] and [95], as implemented on the server at the Bax laboratory. These values were used to calculate conformational chemical shifts (Δδ), as the experimentally measured chemical shift (δ_exp_) minus the calculated chemical shift (δ_coil_). Segments of five or more residues with Δδ^13^Cα > 0.3 ppm and Δδ^13^CO > 0.3 were considered to have a significant preference to form an α-helical segment. When appropriate, families of representative preferred conformers were obtained using the program CYANA 3.98 [96]using the chemical shift data to delimit helical segments. The conformers with the lowest energy functions were chosen to be represented in the figures.

#### Coupling Constants

For segment 5, as an additional, independent test, a 3D HNHA spectrum were recorded and the ratio of the ^1^Hα-^1^HN crosspeak to the ^1^HN-^1^HN diagonal peak intensities in the 3D HNHA spectrum was utilized to calculate the ^3^J_HNCHα_ coupling constants following the procedure of [97]. ^3^J_HNCHα_ coupling constants were also measured for hCPEBpep2 using the 2D ^1^H-^1^H COSY spectrum. Utilizing the Karplus equation [98], these ^3^J_HNCHα_ coupling constants can be related at the backbone ϕ angle, which is different for α-helical, statistical coil and β-strands.

#### Relaxation

To assess the dynamics on the ps–ns time scales, the heteronuclear ^15^N{^1^H} NOE (hNOE) of backbone amide groups was registered as the ratio of spectra recorded with and without saturation in an interleaved mode. Long recycling delays of 13 s were used. Two sets of experiments, one at 25 °C and one at 5 °C were recorded at 800 MHz. Uncertainties in peak integrals were determined from the standard deviation of intensities from spectral regions devoid of signal which contain only noise.

In addition, R_1_ρ relaxation rates, which are sensitive to the presence of preferred, rigid conformers on slower μs-ms timescales were measured by recording two sets of ten ^1^H-^15^N correlation spectra with relaxation delays at 8, 300, 36, 76, 900, 100, 500, 156, 200 and 700 ms. One set of experiments was recorded at 25 °C and the second was recorded at 5 °C. The relaxation rates were calculated by least-squares fitting of an exponential decay function to the data using NMRPipe [99]. As an additional check, the data were also analyzed independently by using the program DynamicsCenter 2.5.2 (Bruker Biospin).

**^13^C detected relaxation experiments** were measured for segment 1 to confirm the presence of a rigid N-terminus and to determine proline residue imino ^15^N relaxation rates. Transverse relaxation rates (R_2_) of hCPEB3 segment 1 imino ^15^N nuclei were measured by a ^13^C-detected c_hcacon_nt2_ia3d pulse sequence [87] as a pseudo 3D experiment time composed of nine 2D experiments with relaxation delays of 15.9, 79.2, 158.4, 269.3, 396.0, 554.4, 712.8, 871.2 and 1030 milliseconds over a ^15^N chemical shift range that is selective for the Pro ^15^N chemical shifts (132 – 140 ppm). This experiment, and a similar one with a wider sweep width to also measure the ^15^N T_2_ relaxation of all 20 imin/amino-acid residues, were recorded at 25 °C, without non-uniform sampling or linear prediction. Following Fourier transformation, IPAP virtual decoupling and baseline correction, the peaks were integrated with TOPSPIN 4.0.8 or alternatively NMRPipe by a different operator for comparison. A single exponential decay curve was then fit to the peak integral versus time data to calculate R_2_ rates for each position.

## Acknowledgments

NMR experiments were performed in the “Manuel Rico” NMR Laboratory (LMR) of the Spanish National Research Council (CSIC), a node of the Spanish Large-Scale National Facility (ICTS R-LRB). We are grateful to Dr. Albert Galera-Prat for help in the initial stages of the study, and to Dr. Miguel Mompeán and Dr. Javier Oroz for critical comments on the manuscript. This study was supported by projects SAF2016-76678-C2-1-R (MC-V) and SAF2016-76678-C2-2-R (DVL) from the Spanish Ministry of Economy and Competitivity and PID 2019-109306RB-I00/AEI/10.13039/501100011033 from the Spanish Ministry of Science and Innovation (DVL). The authors declare no competing interests. All data needed to evaluate the conclusions in the paper are present in the paper and/or the Supplementary Materials.

## Author contributions

1. Funding: MC-V & DVL; 2. Research planning & design: DRdM, RH, DP-U, MC-V and DVL. 3. Cloning, sample production and purification: DRdM & RH; 4. Data acquisition: DP-U & DVL; 5. Data analysis: DP-U, DRdM & DVL; 6. Writing: DRdM, RH & DVL. All authors carefully read the MS, provided corrections and approved the final version of the MS.

## Figures and Tables

**Sup. Table 1:**
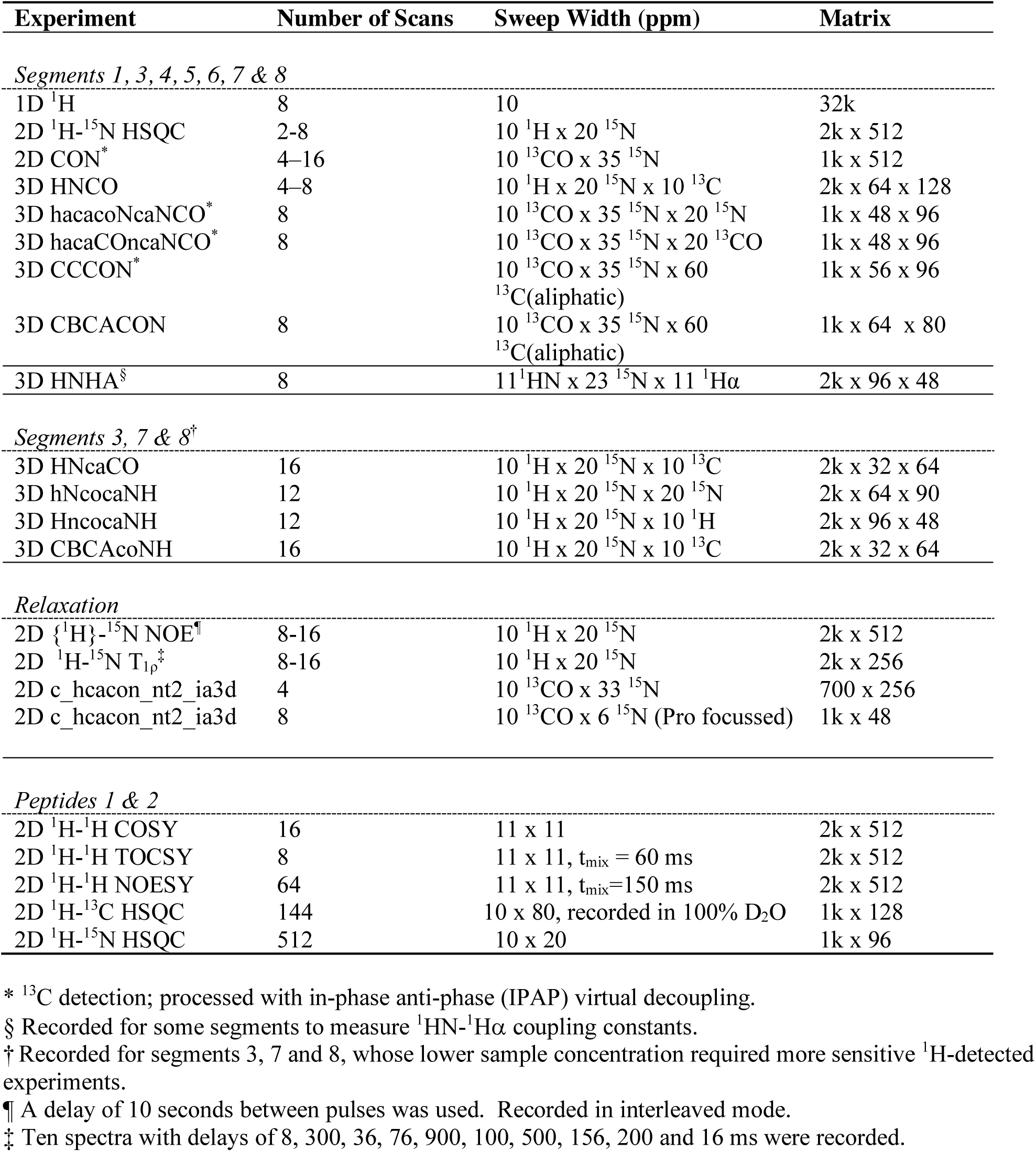
NMR Spectral Parameters.

**Sup. Fig. 1.**
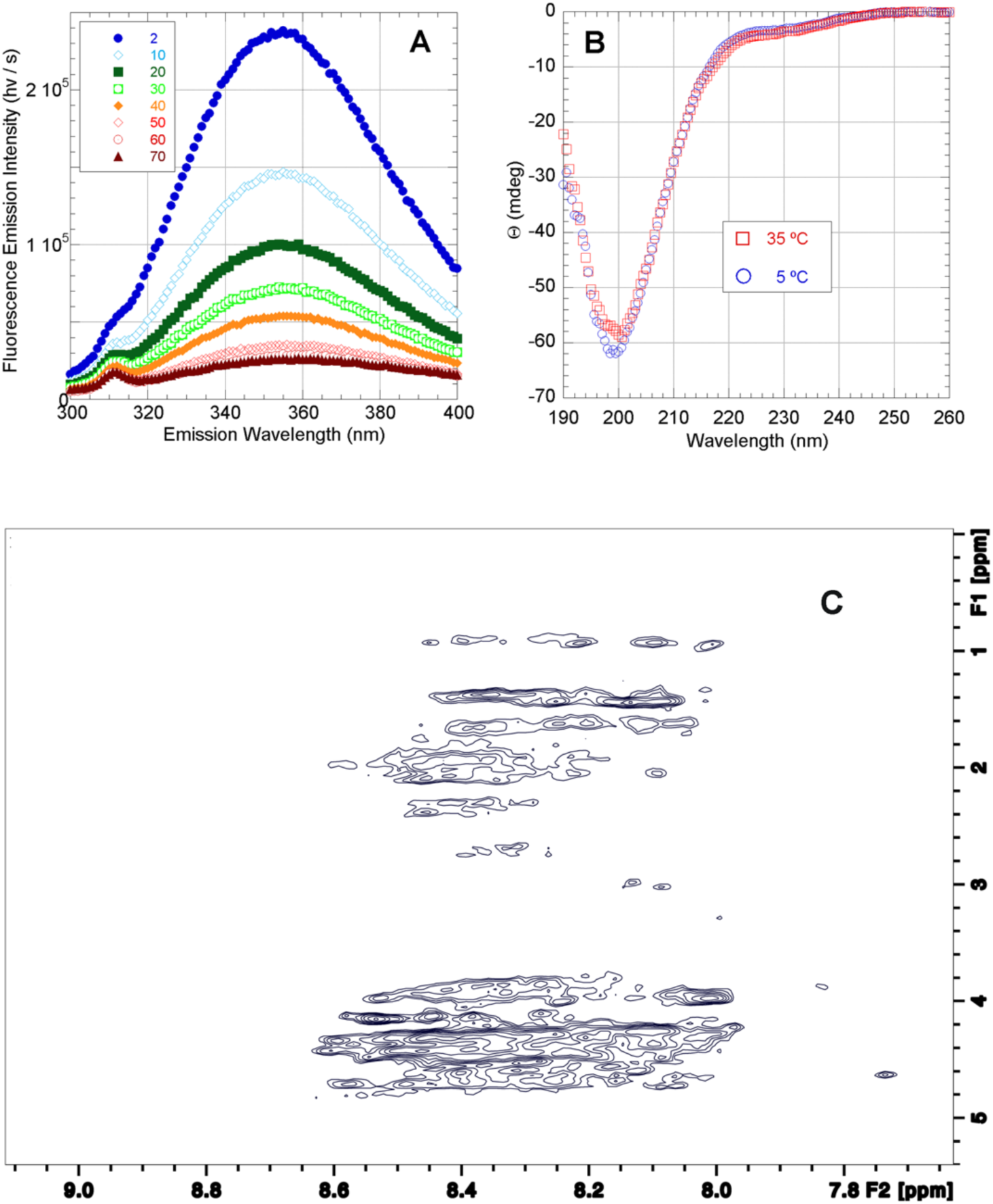
Biophysical Characterization of the Complete hCPEB3 IDR **A.** Fluorescence emission spectra of the complete IDR of hCPEB3 at pH 4.0 in 1.0 mM deuterated acetic acid buffer over temperatures ranging from 2 °C to 70 °C as indicated. The emission maximum over 350 nm is indicative of solvent exposed Trp side chains. This is consistent with the lack of a well packed hydrophobic core. **B.** Far UV-CD spectra of the IDR of hCPEB3 at 5 °C (blue open circles) and 35 °C (red open squares). The observed minimum near 200 nm and the lack of minima near 208 nm, 218 nm and 222 nm and the lack of a maximum at 195 nm are all characteristic spectral features of a statistical coil. **C.** 2D ^1^H-^1^H NOESY spectrum of the hCPEB3 IDR recorded at 25 °C showing the ^1^HN crosspeak region. The small chemical shift dispersion in ^1^HN (8.6 to 7.9 ppm) is a typical feature of disordered and unfolded proteins (López-Alonso *et al.,* 2010) [27].

**Sup. Fig. 2:**
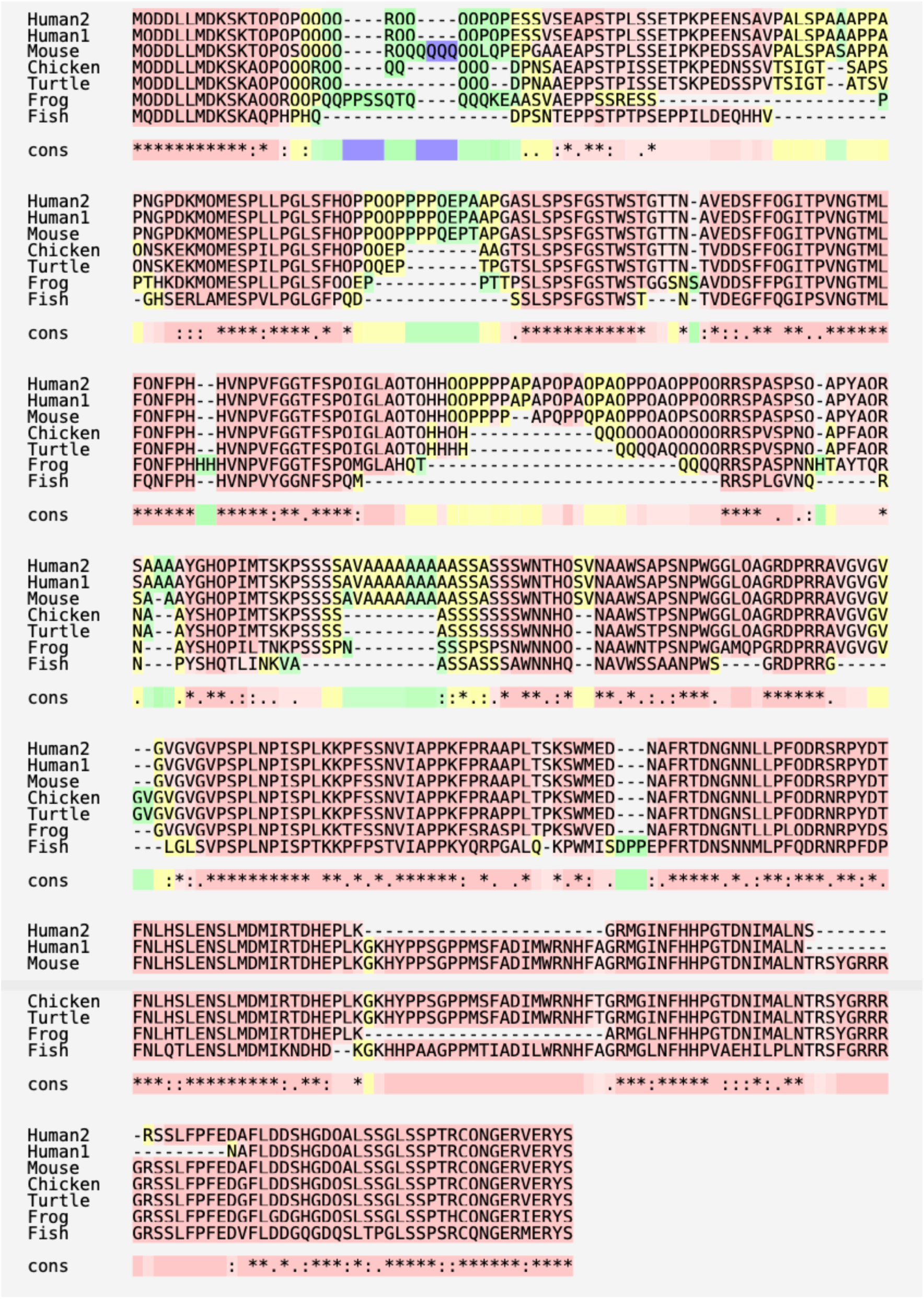
Sequence Alignments of CPEB3 from Representative Vertebrates Results obtained using T-coffee, version 11 (Di Tommaso *et al.,* 2011) [81] using default settings. Sequence is highlighted using a red to blue color scale for highly to poorly conserved residues, respectively. Note that because of the small space between lines, some “Q”s appear as “O”s. “Cons” indicates the level of sequence conservation: “*” = strictly conserved, “:” = well conserved, “.” moderately conserved.

**Sup. Fig. 3.**
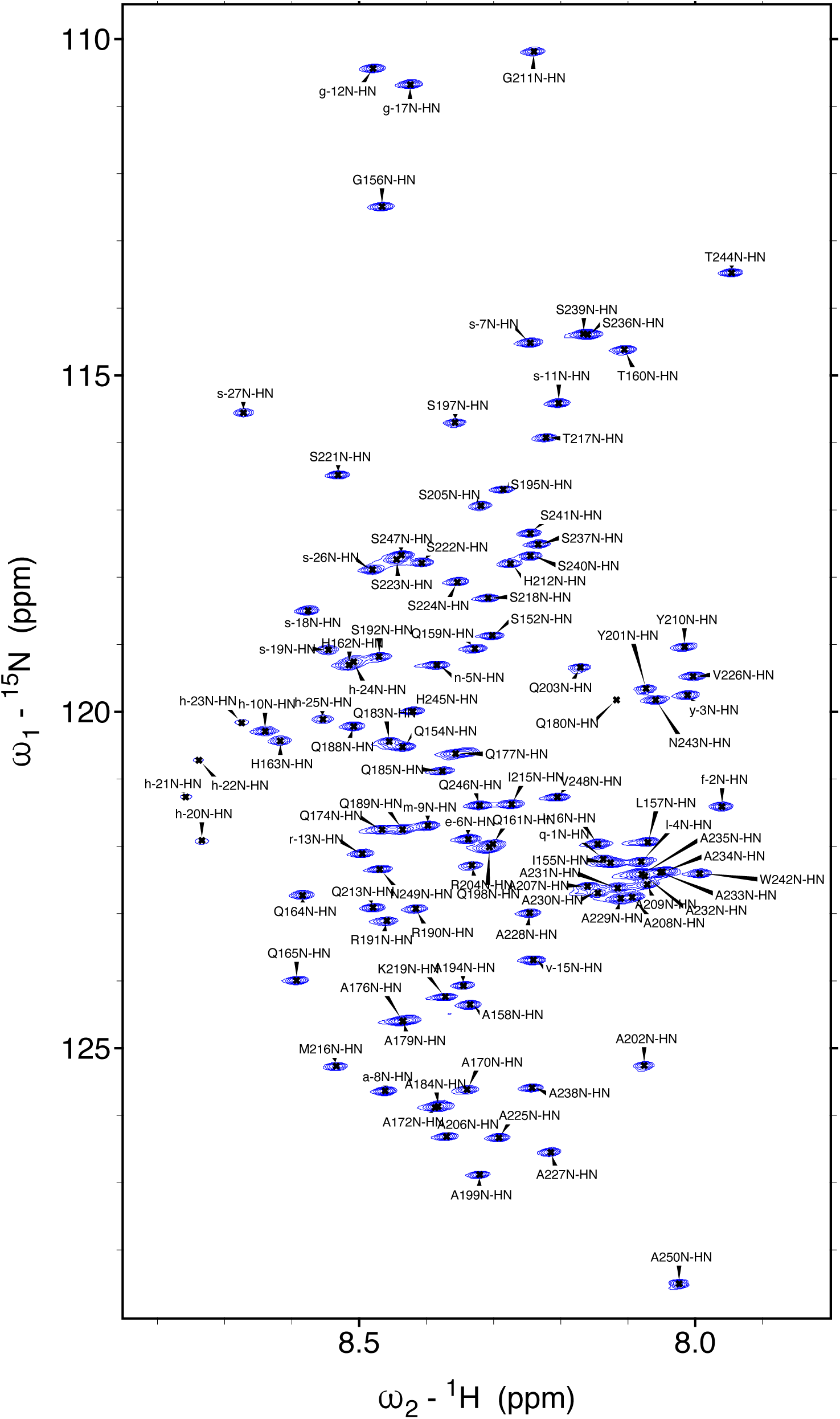
2D ^1^H-^15^N HSQC NMR Spectrum of hCPEB3, Segment 4 ^1^H-^15^N correlations were recorded at 25°C in 1 mM acetic acid, pH 4. Note the cluster of Ala signals in belonging to the polyA stretches near 8.05 ppm ^1^H and 122.5 ppm ^15^N. Their position contrasts with those of the isolated Ala residues, whose δ ^15^N > 125 ppm. Signals labeled in lower case with negative numbers correspond the His/TEV tag: M_-29_GSSHHHHHHSSGLVPRGSHMASENLYFQ_-1_

**Sup. Fig. 4.**
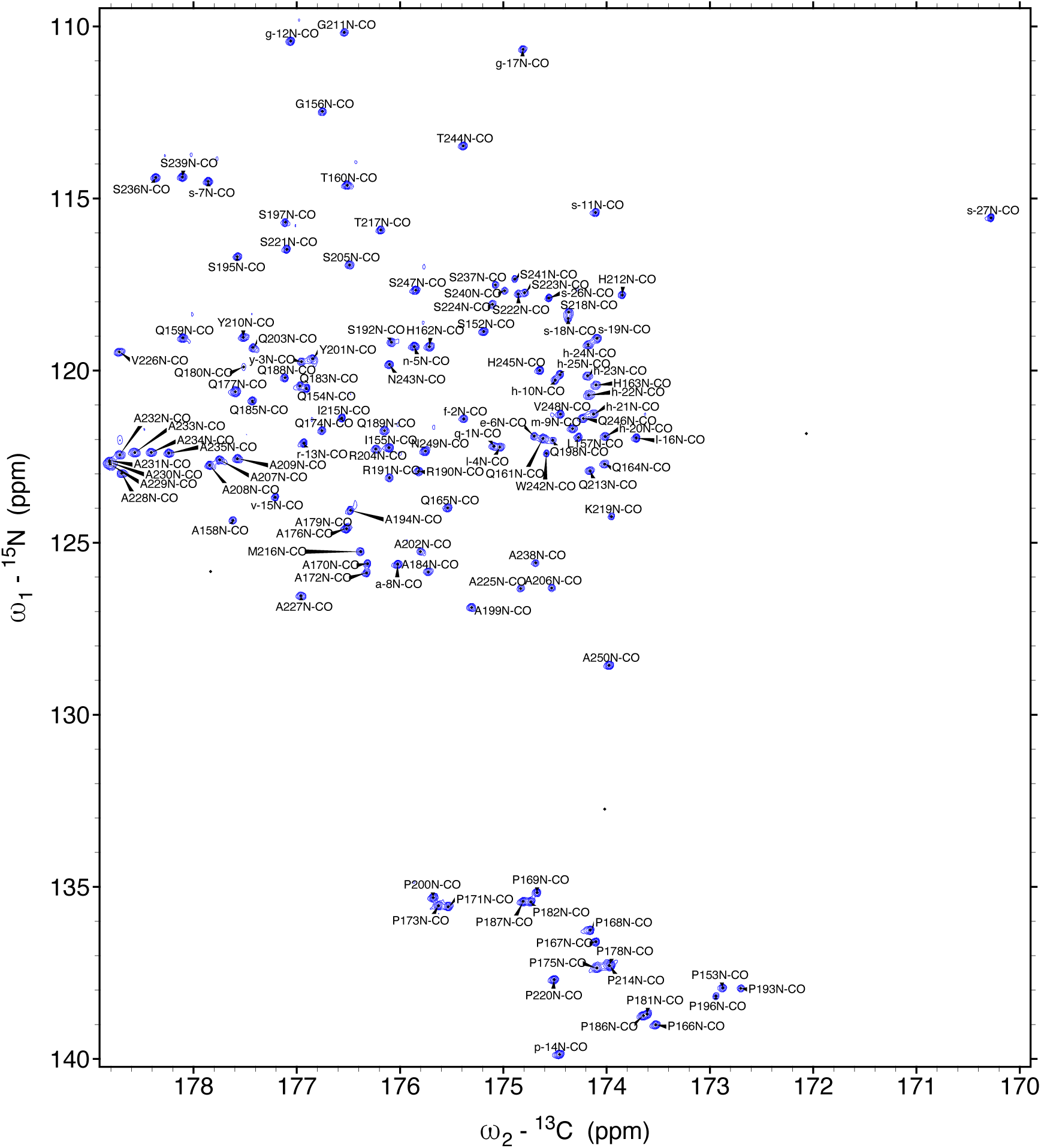
2D ^13^CO-^15^N NMR Spectrum of hCPEB3 ^13^CO-^15^N correlations were recorded at 25°C in 1 mM acetic acid, pH 4. Here, the signals of the long polyAla stretch, which appears around 122.5 ppm for ^15^N x 178.5 ppm for ^13^CO shows superior resolution relative to the ^1^H-^15^N HSQC spectrum. Their position at higher ^13^CO and lower ^15^N chemical shift values relative to isolated Ala and to coil values indicates an alpha helical conformation. For simplicity, the label refers to the ^13^CO of the *i-1* residue. Signals labeled in lower case with negative numbers correspond the His/TEV tag: M_-29_GSSHHHHHHSSGLVPRGSHMASENLYFQ_-1_

**Sup. Fig. 5.**
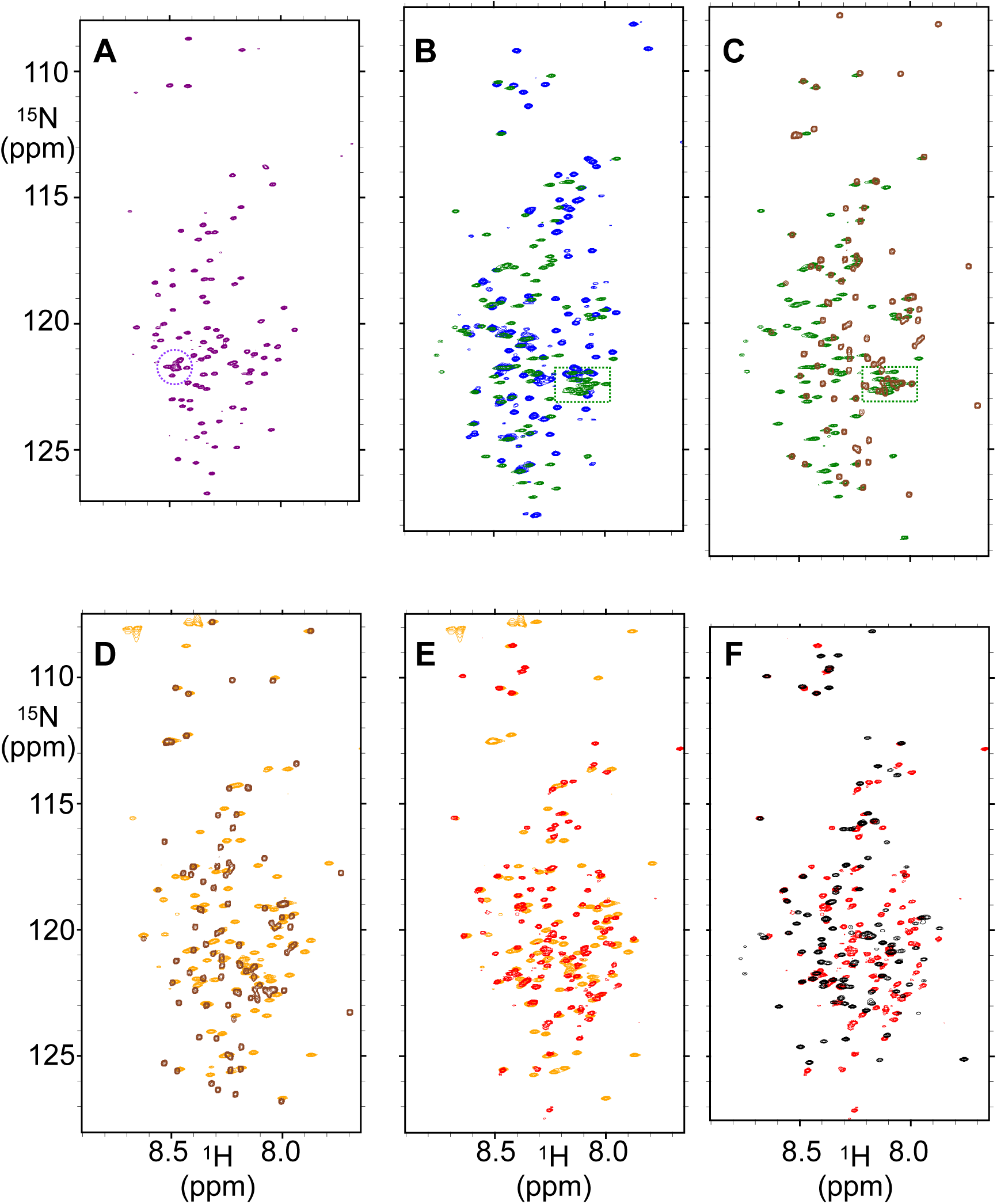
^1^H-^15^N HSQC spectra of hCPEB3 IDR Segments 2D ^1^H-^15^N HSQC spectra of hCPEB3 IDR recorded at pH 4, 25°C on: A. Segment 1 (**purple**), signals from the polyQ rich segment are circled. B. Segments 3 (**blue**) and 4 (**green**). Signals from the polyA rich segment are boxed. C. Segments 4 (**green**) and 5 (**brown**). Signals from the polyA rich segment are boxed. D. Segments 5 (**brown**) and 6 (**orange**). E. Segments 6 (**orange**) and 7 (**red**). F. Segments 7 (**red**) and 8 (**black**)

**Sup. Fig. 6.**
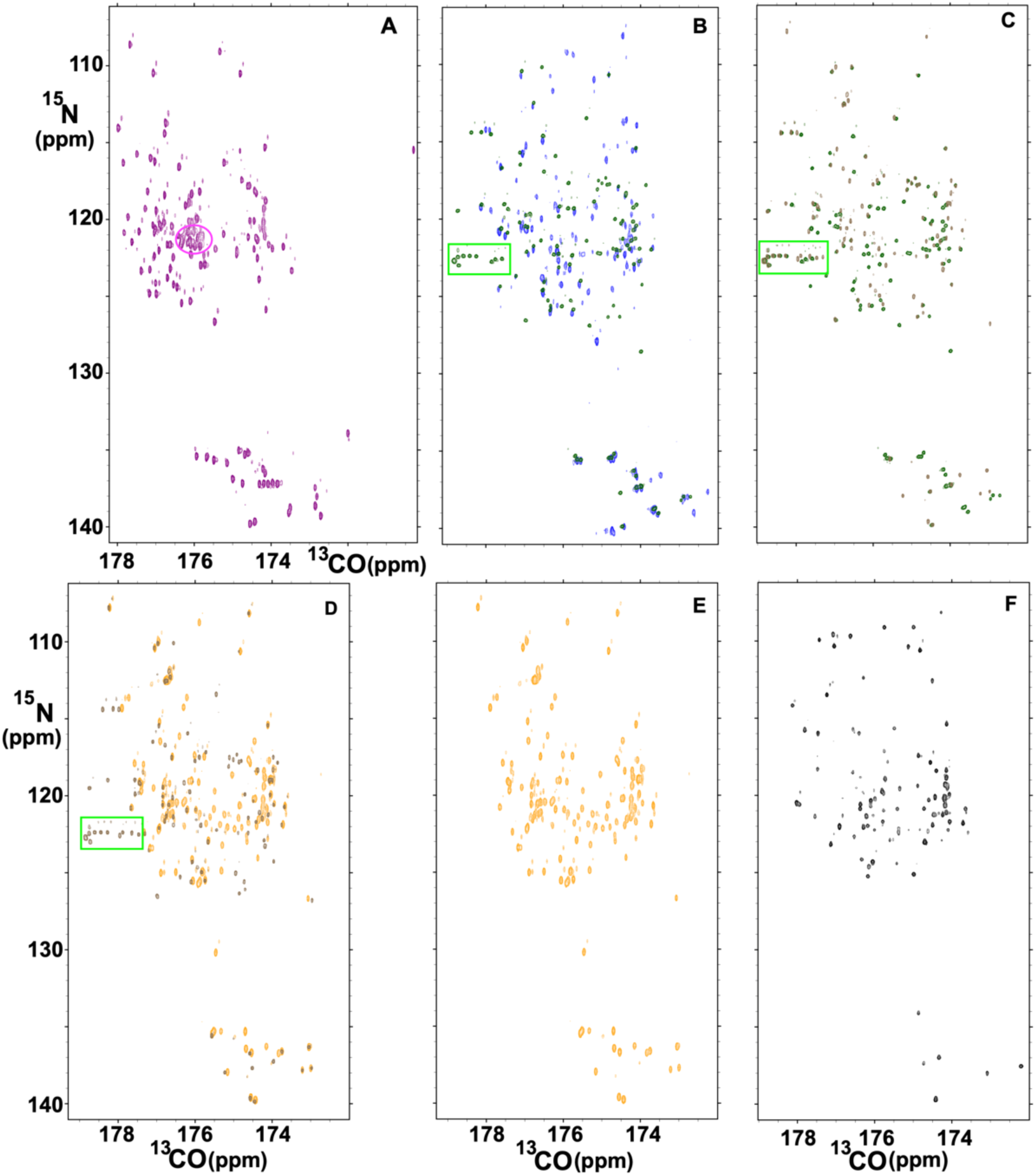
2D ^13^CO-^15^N spectra of hCPEB3 IDR Segments 2D ^13^CO-^15^N HSQC spectra of hCPEB3 IDR recorded at pH 4, 25°C on: A. Segment 1 (**purple**), signals from the polyQ rich segment are circled in magenta. B. Segments 3 (**blue**) and 4 (**green**). Signals from the polyA rich segment are boxed in **light green**. C. Segments 4 (**green**) and 5 (**brown**). Signals from the polyA rich segment are boxed in **light green**. D. Segments 5 (**brown**) and 6 (**orange**). E. Segment 6 (**orange**). F. Segment 8 (**black**) The spectrum of segment 1 is displaced because its optimal spectral range was distinct relative to the other spectra. The weak satellite peaks appearing to the right and above the main peaks arise from deuteration.

**Sup. Fig. 7.**
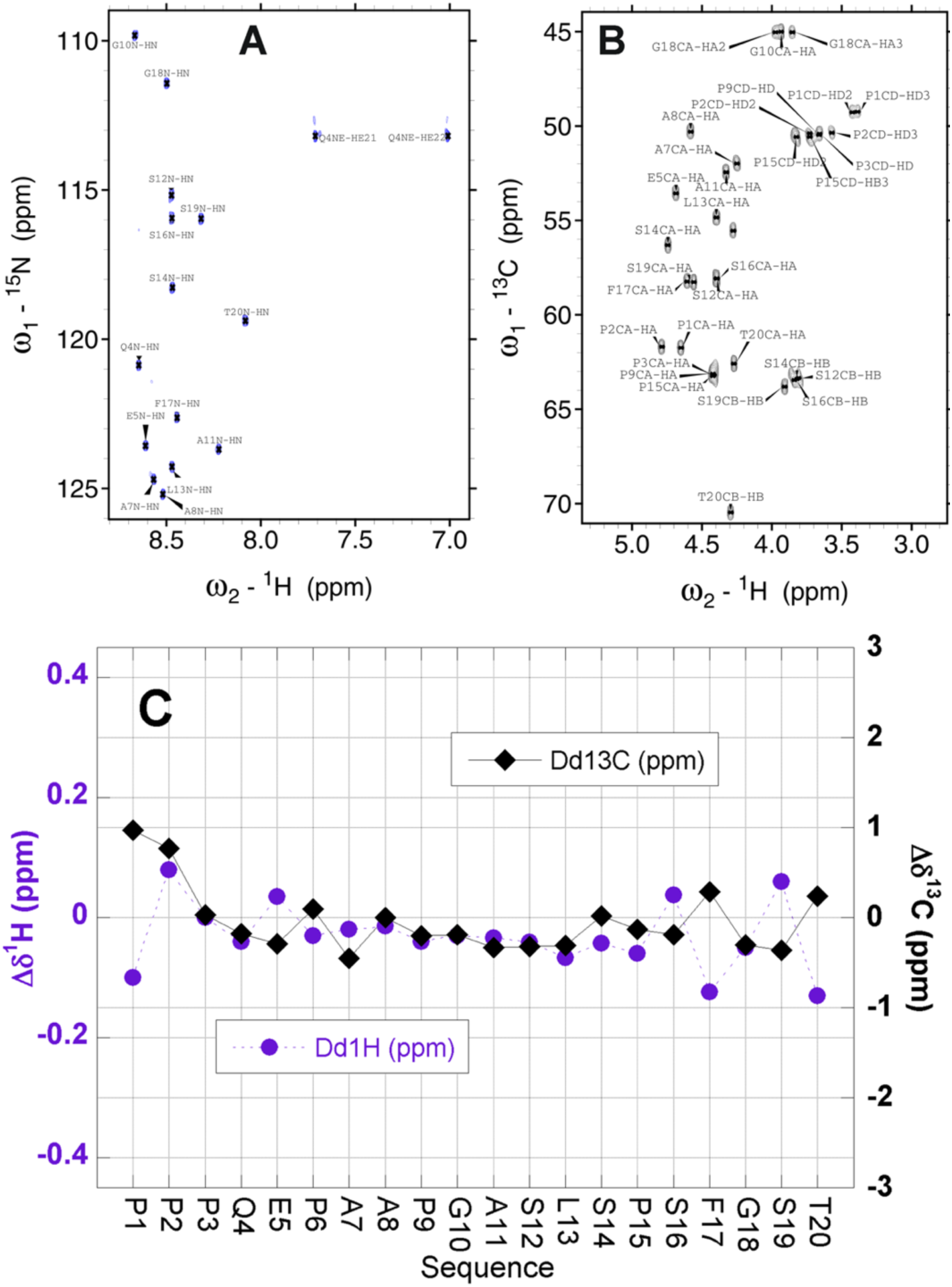
Corroboration of small to negligible populations of α-helix or β-strand conformations in residues 91 - 110 of hCPEB3. **A.** The assigned ^1^H-^15^N HSQC spectrum of a peptide corresponding to residues 91 - 110 of hCPEB3. The low chemical shift dispersion in the ^1^HN dimension is a hallmark of a statistical coil. The residue number labels correspond to this peptide, to calculate the residue number in the full length protein, add 90. **B.** The ^1^H-^13^C HSQC spectrum of the same peptide, showing the assigned ^1^Hα-^13^Cα, ^1^Hδ-^13^Cδ Pro and ^1^Hβ- ^13^Cβ Ser & Thr correlations. **C.** Conformational chemical shifts of ^1^Hα (left y-axis, **purple circles**) and ^13^Cα (right y-axis, **black diamonds**). Values of -0.41 ppm (^1^H) and +3.1 ppm (^13^C) correspond to 100% α-helix and values of +0.41 (^1^H) and -3.1 ppm (^13^C) correspond to 100% β-strand. Here residues show small conformation chemical shifts except for N- and C-termini, whose values are perturbed by end effects, and residues near Phe17 (Phe 107 in the full length protein) whose values are altered by ring current effects.

**Sup. Fig. 8.**
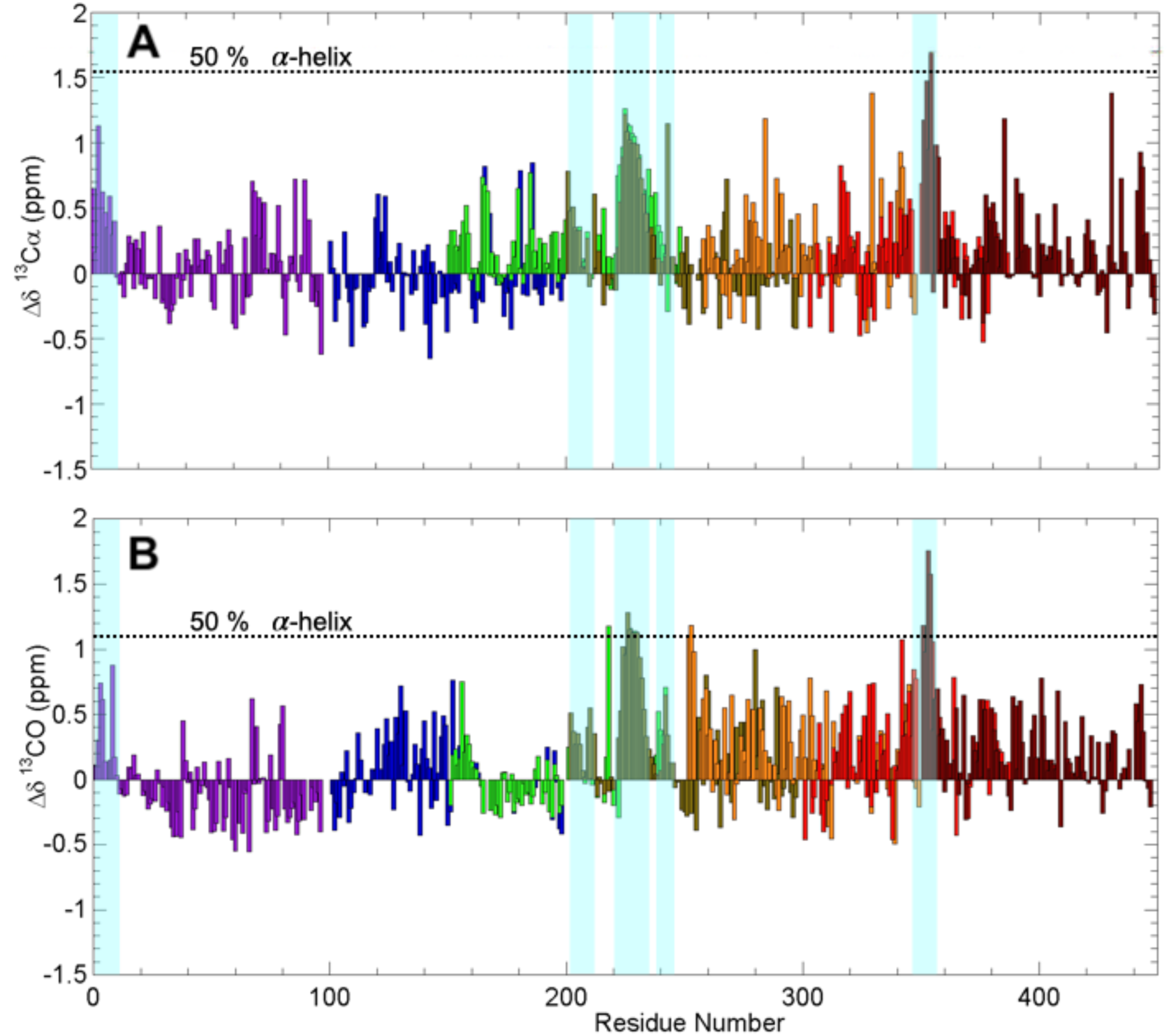
Conformational chemical shifts reveal the presence of partially populated secondary structure in the hCPEB3 IDR. Conformational chemical shifts (Δδ) at 25°C, calculated as the experimentally measured chemical shift (δ_exp_) minus the chemical shift expected for statistical coil (δ_coil_) for ^13^Cα (*top panel*) and ^13^CO (*bottom panel*) under these conditions. Data are colored according to segment: 1=**purple**, 3=**blue**, 4=**green**, 5=**brown**, 6=**orange**, 7=**red**, 8=**maroon**. Negative values Δδ values of ^13^Cα and ^13^CO are characteristic of extended conformations like β-strands. Values of +1.55 ppm for Δδ ^13^Cα and +1.1 ppm Δδ ^13^CO are expected for 50% α-helix; these values represented as dotted lines. Stretches of positive Δδ for ^13^Cα and Δδ ^13^CΟ values which evince partial α-helix formation span residues 1-10, 202-210, 222-234, 238-246 and 346-356 and are shaded cyan.

**Sup. Fig. 9.**
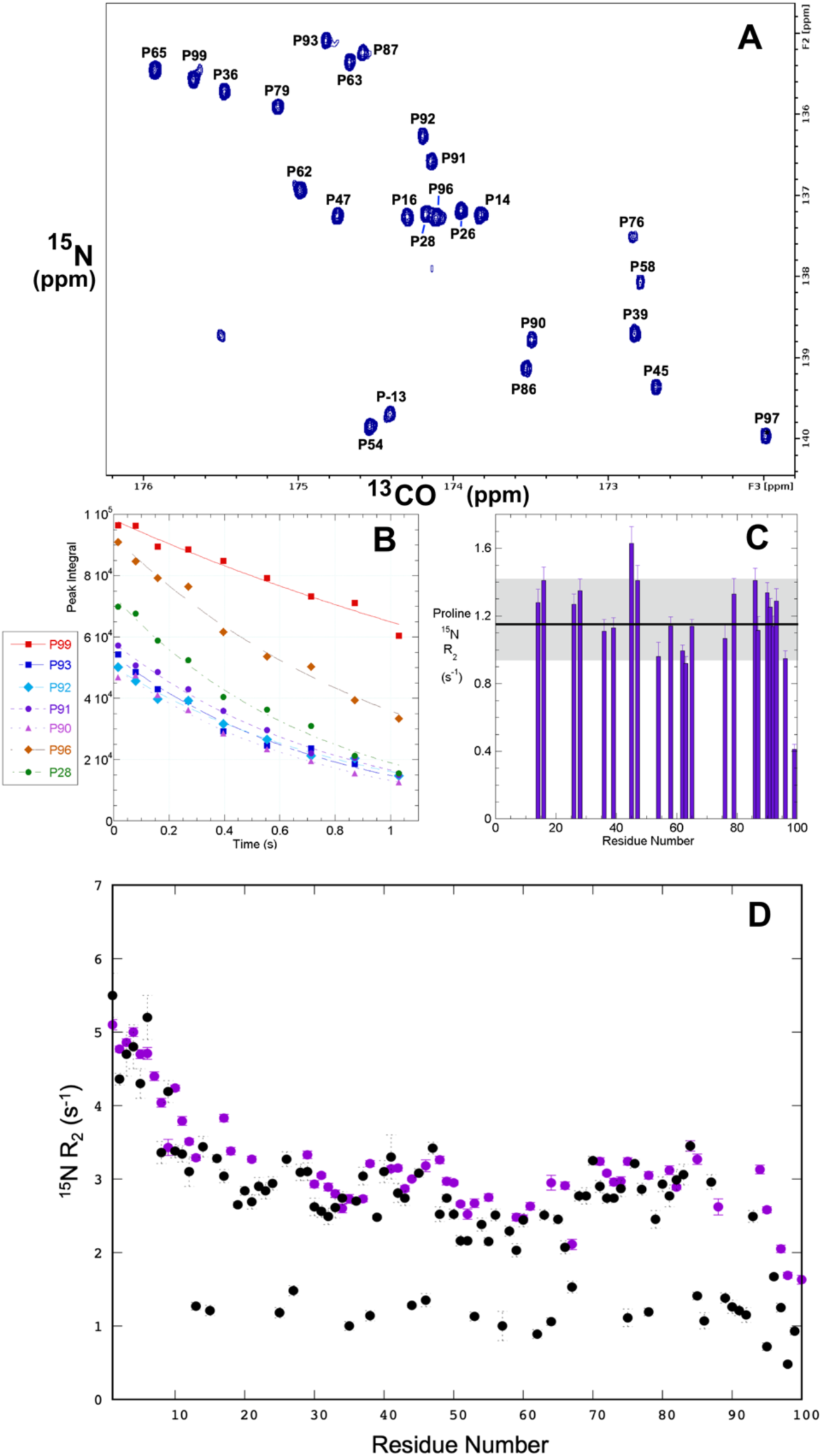
^13^C-detected ^15^N Relaxation Experiments of hCPEB3 segment 1. **A.** The ^13^C-detected ^15^N relaxation experiment obtained using a 15.9 ms delay. Proline ^15^N atoms are labeled. **B**. Fits of some represantative proline ^15^N nuclei are shown Peaks “P-13”, which comes from the His tag, and P97, whose resonance is folded, were not included in the analysis. The R_2_ rates and their one standard deviation uncertainties of their fits (error bars) from the Topspin analysis are shown in **panel C**. NMRPipe analysis yielded similar results (data not shown). The black line marks the mean R_2_ rate (1.18 s^-1^) and the gray shaded area is the one standard deviation (+/- 0.24 s^-1^) from the mean of all the proline R_2_ rates. As previously observed in two recent studies (Murrali *et al.,* (2018); Mateos *et al.,* 2020), the R_2_ rates for the proline imine ^15^N are consistent with *trans* Xaa-Pro peptide bond (Mateos *et al.,* 2020) and are significantly slower relative those for the 19 amino acid residues. The residues near the C-terminus, especially P99, shows slower relaxation which is indicative of high flexibility (**panel C**). In contrast, the prolines of the two “PQP” mini-motifs (P14 & P16) and (P26 & P28) flanking the Q_4_RQ_4_ motif show slightly higher R_2_ rates as do P45 & P47 (which are found within a cluster of charged residues) and P90, P91, P92 & P93 (which adopt a short PPII helix). This is indicative of slightly less flexibility. Similar trends were observed recently in isolated Pro residues and a run of four consecuetive proline residues in the ID4 linker domain of the CREB binding protein (Murrali *et al.,* (2018). Finally the ^13^C-detected relaxation experiment was repeated for all hCPEB3 segment 1 residues using a broad sweep width and the results are shown in **panel D** as black circles along the results obtained using a ^15^N detected experiment, which are colored purple. The good agreement is a further confirmation of the presence of a rigid, partially populated α-helix at the N-terminus of Segment 1 (**Figure 2, Sup. Fig. 10 & 12**).

**Sup. Fig. 10.**
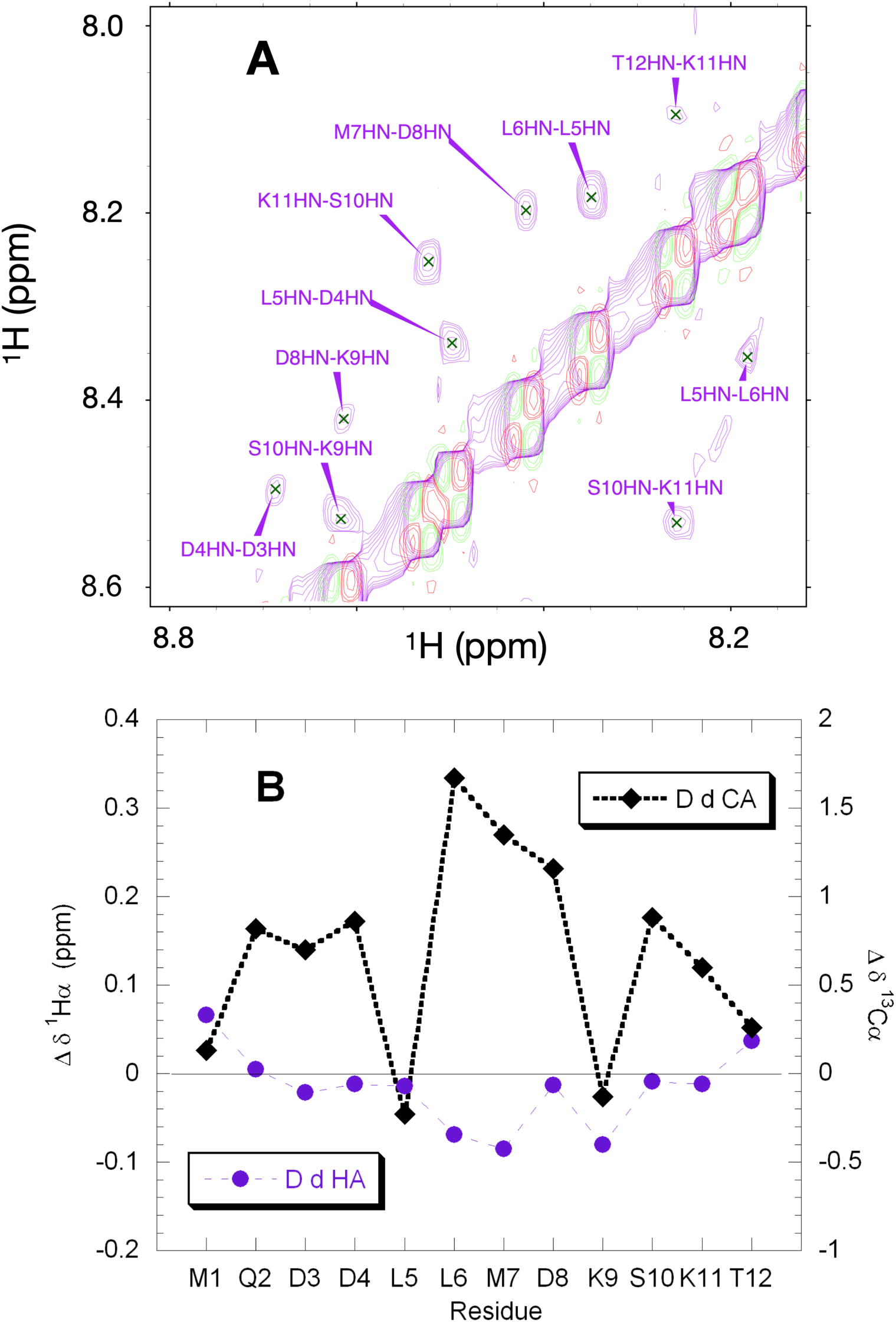
Partial Formation of an α-Helix in the N-terminal Residues of hCPEB3 **A.** 2D ^1^H-^1^H NOESY (80 ms mixing time, **purple**) of a peptide corresponding to the initial residues of hCPEB3. Sequential NOE correlations, which are consistent with α-helical structure, are labeled. **Red** and **green** peaks along the diagonal are from the 2D ^1^H-^1^H COSY spectrum. **B.** Conformational ^1^H (**purple**) and ^13^C (**black**) chemical shifts at 5 °C for a peptide corresponding to the initial residues of hCPEB3 20% hexafluoroisopropanol (CF_3_-CHOH-CF_3_) The positive Δδ^13^C and negative Δδ ^1^H values are indicative of the formation of significant α-helix formation.

**Sup. Fig. 11.**
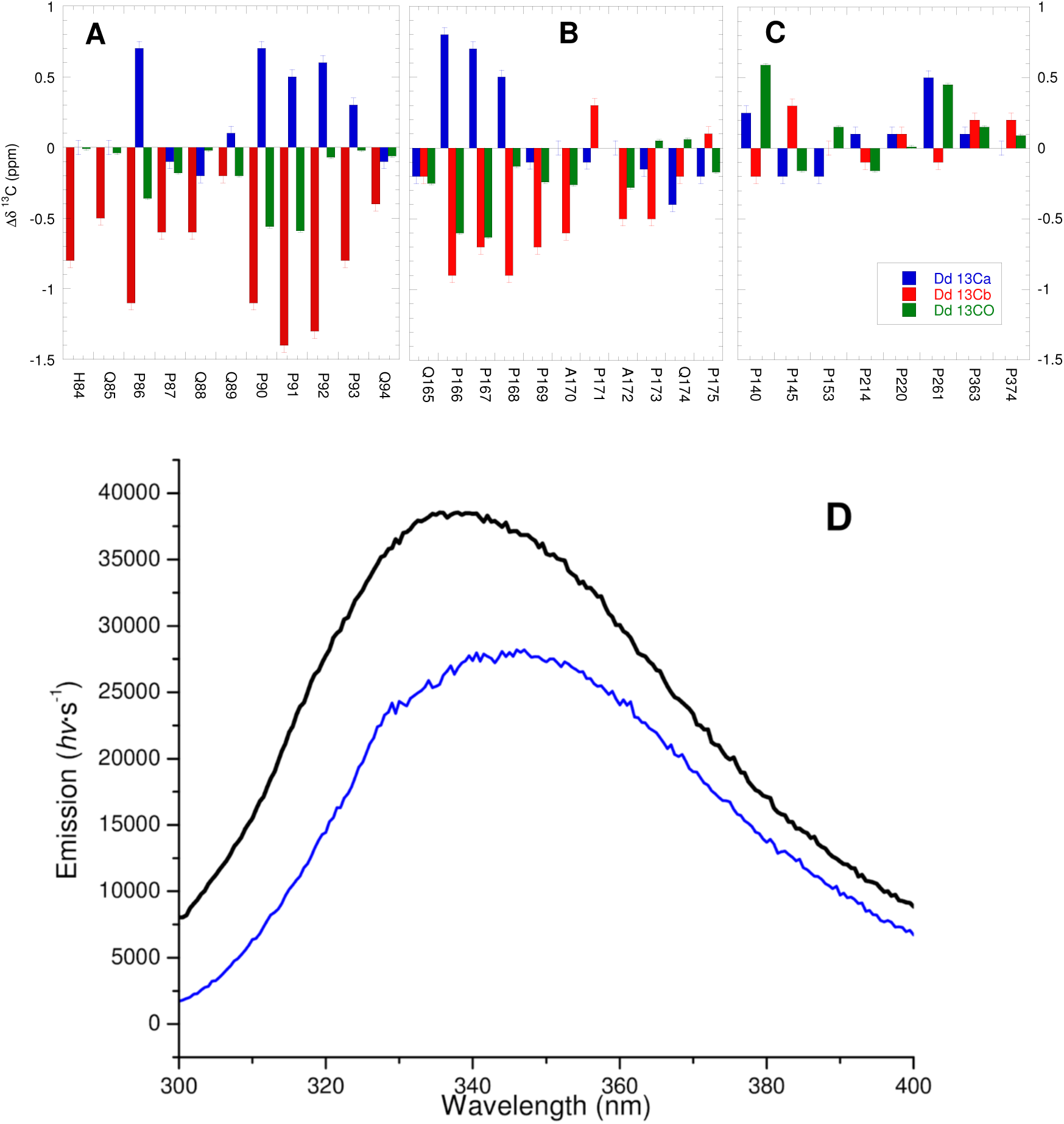
The Consecutive Proline Residues of hCPEB3 Show a Characteristic Pattern of Conformational Chemical Shifts and Weak Binding to Profilin. **A**. and **B**. Conformational chemical shifts (Δδ) for ^13^Cα (**blue** bars), ^13^Cβ (**red**) and ^13^CO (**green**) nuclei for residues in two Pro-rich stretches, H84-Q94 and Q165-175, each of which contains four consecutitive Proline residues which will be locked in the polyproline II helical conformation. The Δδ values of residues P90, P91, P92, P168, P167 and P168 were averaged to obtain the values given in Table 3 on the main text. **C.** Isolated proline residues, *i.e.* those which do not have another proline residue within two anterior or posterior positions along the sequence, do not show a pattern of significant Δδ deviations. **D.** Fluorescence emission spectra of 8.3 µM human profilin 1 in the absence (blue spectrum) or presence (black) of 4.5 mM acGPPPPAPAPQPam peptide at 20 °C in 10 mM K_2_HPO_4_, 75 mM KCl, (pH 8.0) at 20 °C. The enhanced, blue-shifted emission is indicative of binding [34].

**Sup. Fig. 12.**
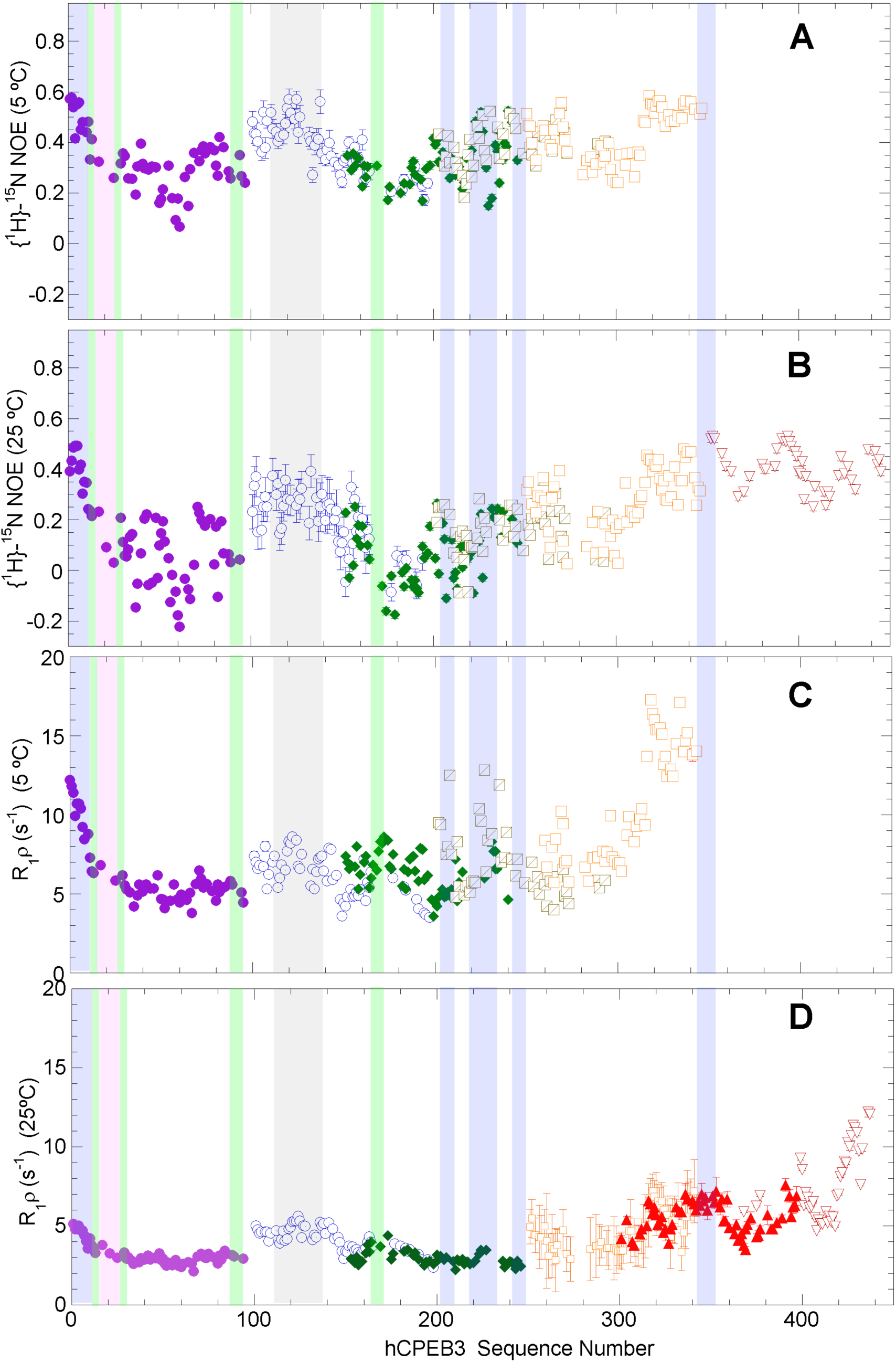
Residue Level Dynamics of hCPEB3’s Instrinsically Disordered Region Residue-level dynamics on ns/ps time scales at 5°C (**A**) and 25°C (**B**) and on μs/ms at 5°C (**C**) and 25 °C (**D**). In panels **A** and **B**, a value of 0.86 is expected for completely rigid H-N groups; negative values are characteristic of a high flexibility. In panels **C** and **D**, while values less than 4 s^-1^ are hallmarks of flexibility, higher values mean decreased mobility. Data from individual segments are colored differently: segment 1 = **purple filled circles**, segment 3 = **blue open circles**, segment 4 = **green diamonds**, segment 5 = **brown barred open squares**, segment 6 = **orange open squares**, segment 7 = **filled red triangles**, segment 8 = **open inverted maroon triangles**. No data are represented for segments 7 and 8 at 5°C, segment 5 in panel **D**, and segment 7 in panel **B** as the spectra obtained were of insufficient quality. The lack of other values is due to ^1^H^15^N signal overlap or proline residues whose nitrogen lacks a hydrogen. Error bars represent uncertainties as estimated from peak intensity signal/noise for panels **A** and **B** and as obtained from the fit of a signal exponential decay function to the peak intensity versus delay time data for panels **C** and **D**. The error bar is frequently smaller than the data symbol. Elements of partial structure/interest are shaded **blue** for α-helices and **gray** for hydrophobic segments; these elements generally show decreased mobility. By contrast, the polyQ tract (shaded **magenta**) is more flexible. Whereas the dynamics of the PPII tracts (**green**) can now be assessed directly as these imine residues lack ^1^HN, nearby residues do show relatively high mobility.

**Sup. Fig. 13.**
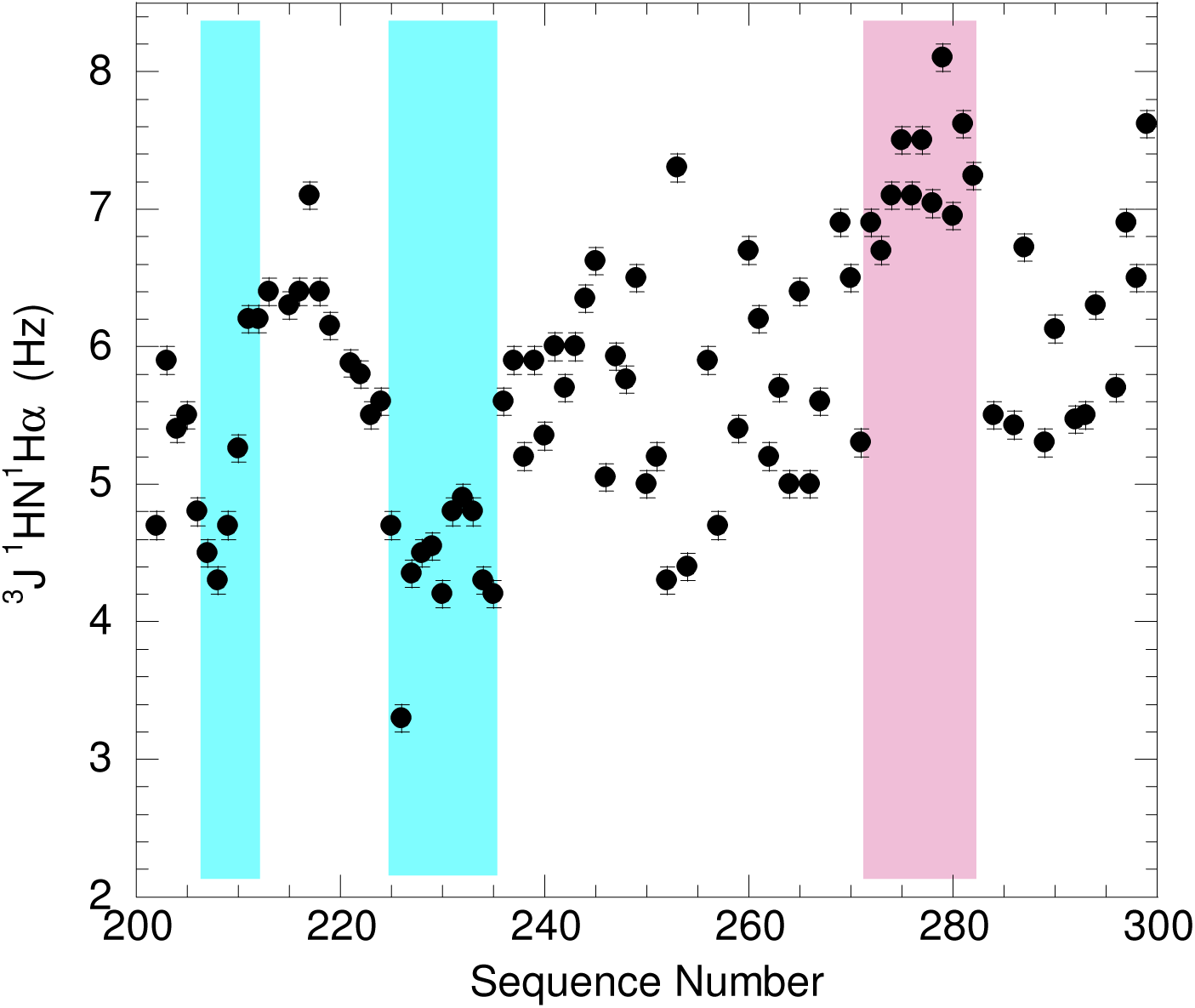
^1^HN-^1^Hα Coupling Constants for Segment 5 Confirm the Presence of α-Helices. Intraresidual three bond ^1^HN-^1^Hα coupling constants (^3^J_1HN1Hα_) for segment 5 of hCPEB3. Regions with ^3^J_1HN1Hα_ < 5 (shaded **cyan**) correspond to two α-helices, in line with the results based on chemical shift deviations (*see* ***Fig. 3*** *in the Main Text*). By contrast, the last, modestly populated α-helix identified by chemical shift deviations (A238-Q246), does not show significantly lower ^3^J_1HN1Hα_ values. A zone with ^3^J_1HN1Hα_ > 7 Hz (shaded **rose**) indicates extended conformations. This zone spans the (VG)_5_ motif.

**Sup. Fig. 14.**
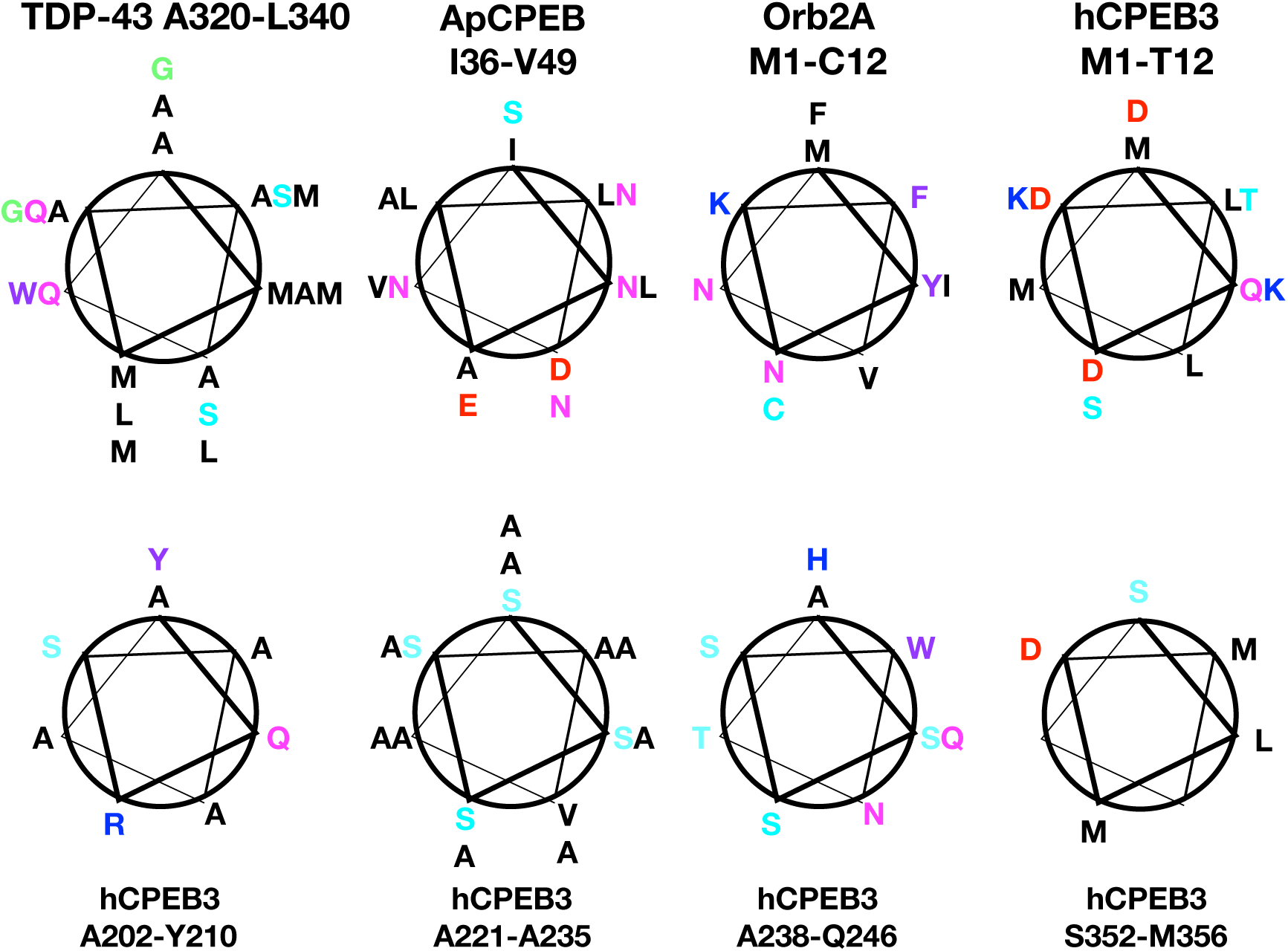
Helices from Pathological and Functional Amyloids Are Stabilized by Distinct Interactions. Helical wheel diagrams of TDP-43, ApCPEB, Orb2A and hCPEB3, nonpolar residues are colored **black**, aromatics=**purple**, Q/N=**magenta**, E/D=**red**, H/K/R=**blue**, G=g**reen**, C,S,T=**cyan**. The five hCPEB3 helices were identified here by NMR data. The TDP-43 helix was identified by NMR by Lim *et al.,* 2016. [51] The ApCPEB and Orb2A helices are putative as they have not yet been confirmed experimentally.

**Sup. Fig. 15.**
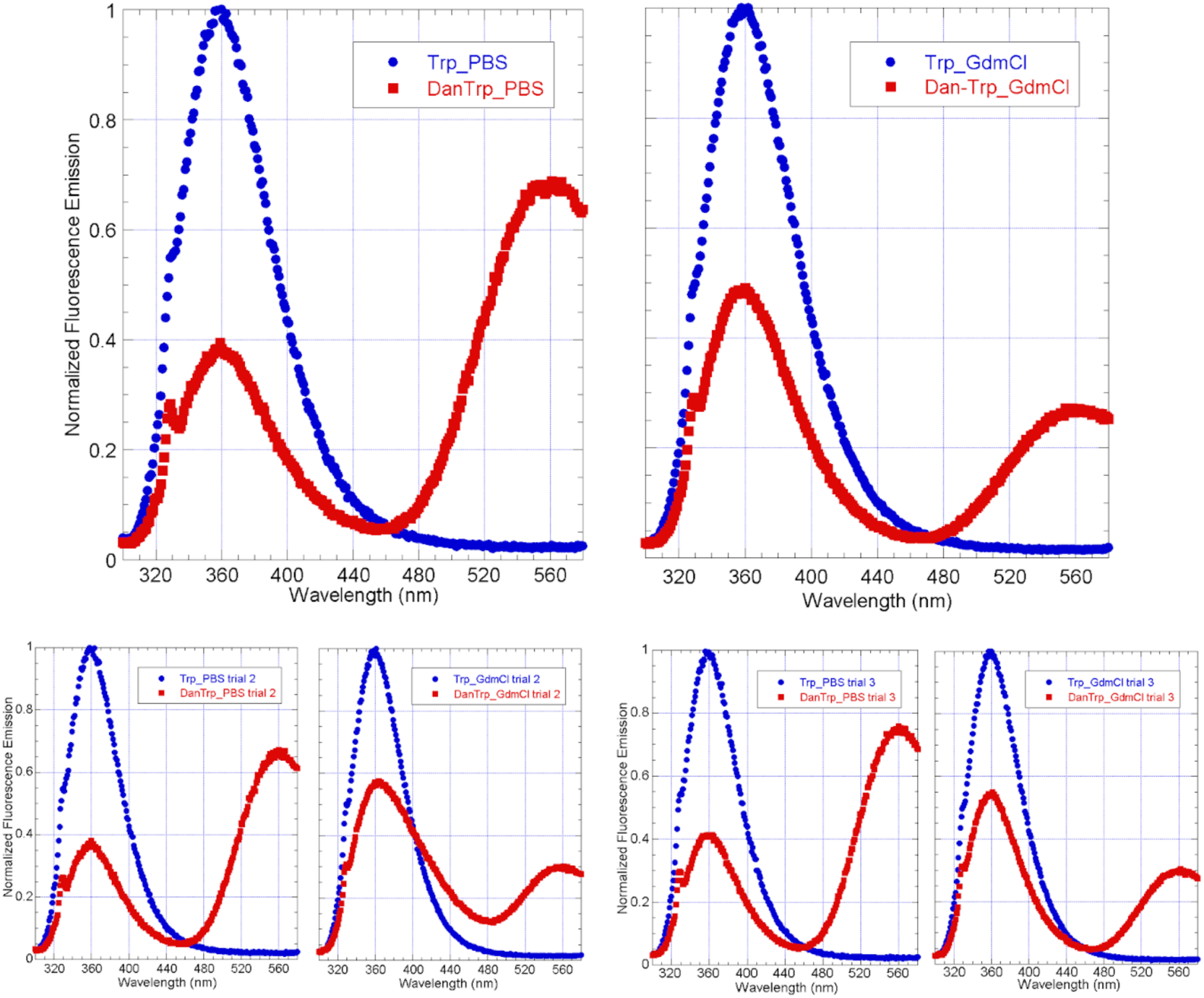
Insight into Interhelix Interactions from FRET Fluorescence spectra of two polypeptides whose concentration was 10.0 µM: MQDDLLMDKSKTGGGGASSSWNTHQ called “Trp” and Dansyl-MQDDLLMDKSKTGGGGASSSWNTHQ called “Dan-Trp”, with residues corresponding to the first and fourth partly populated α-helices of hCPEB3 (shaded pink and cyan, respectively) were recorded. It can be seen in the graph on the *left* (in buffer) that the emission of Trp is high in “Trp” but low in the “Dan-Trp” peptide. This is due to energy transfer of Trp to Dan and thanks to this the Dansyl shines and there is an emission centered at 560 nm which is not seen for the “Trp” peptide. Applying the theory of Förster and making usually assumptions about the rapid movement and averaged orientation of the donor and acceptor groups, and using the known “Förster distance” (R_0_) of 23.6 angstroms Lakowicz *et al.* (1990), the mean distance between the Dansyl and Trp can be calculated to be 22 angstroms, which is significantly lower than the mean value (41 angstroms) expected for a statistical coil of 21 residues (the number of residues separating the Dansyl and Trp groups) as calculated on the basis of the data complied by Fitzkee and Rose (2004) PNAS·USA **101**(34):12497-12502 as well as the results reported by Tanford *et al.* (1966) J. Biol. Chem. **241**(8) 1921-1923. On the other hand, in the presence of 7.4 M GdmCl, in the graphs shown on the *right*, the Trp emission intensity decrease due to Dansyl is smaller, and also the emission intensity of Dansyl is less intense. Two repetitions of the experiment are shown in the lower panels. Based on these results, one can conclude that the Trp and Dansyl groups are farther away in the presence of the denaturant, due to the unfolding of a more compact structure which is present in the absence of GdmCl. Thus, these new results support the possibility of interhelical contacts in hCPEB3.

